# A Single-Nucleus Transcriptomic Atlas Reveals Cell Type-Specific Responses to OsHV-1 Infection in the Pacific Oyster

**DOI:** 10.64898/2026.05.15.723513

**Authors:** Pooran S. Dewari, Tim Regan, Ambre F. Chapuis, Alexandra Florea, James J. Furniss, Thomas C. Clark, Richard S. Taylor, Tim P. Bean

**Affiliations:** The Roslin Institute, The University of Edinburgh, Easter Bush Campus, Midlothian, EH25 9RG, UK

**Keywords:** Single-nucleus RNA sequencing, Pacific oyster, mollusc, OsHV-1, herpesvirus infection

## Abstract

**Background:** The Pacific oyster (*Crassostrea/Magallana gigas*) is increasingly recognised as a model marine invertebrate. Valued for both ecological and commercial importance, Pacific oysters are farmed widely, supporting global food security by providing a sustainable nutrient-rich source of protein. Despite the significant and recurring economic losses caused by Ostreid herpesvirus (OsHV-1) outbreaks, only a limited number of studies have examined host-pathogen interplay at single-cell resolution. The few available studies largely focus on circulating immune cells (haemocytes), thereby overlooking the complexity of host responses across different tissues and organs.

**Results:** We present a detailed single-nucleus transcriptomic atlas of the whole Pacific oysters, including during OsHV-1 infection. A total of 18 distinct transcriptomic clusters were resolved, capturing major cell populations from the gill, mantle, hepatopancreas, adductor muscle, and haemocytes. Notably, three populations- gill ciliary cells, hepatopancreas cells, and an immune-enriched cluster 1- exhibited pronounced transcriptomic responses to OsHV-1 infection. Across the 6, 24, 72, and 96 hours post-infection (hpi) time course, viral transcripts were detected almost exclusively at 72 hpi, with enrichment primarily in adductor muscle cells and two immune cell populations- immature haemocytes, and hyalinocytes.

**Conclusions:** Our findings suggest potential entry portals and tissue-specific replication sites for the OsHV-1 virus in Pacific oysters. This atlas resource provides a high-resolution cellular framework for understanding host-virus interactions and establishes a foundation for future investigations into herpesvirus pathogenesis in marine invertebrates.

## Background

Pacific oysters, known as *Crassostrea gigas* or *Magallana gigas*, are one of the most extensively farmed aquatic species globally [1]. Originating from the west Pacific Ocean and consumed across East Asian countries for thousands of years, their rapid growth rates, high tolerance of variable environmental conditions, amenability to hatchery production, and resistance to many common pathogens of native species, have resulted in uptake across the globe. They are now farmed heavily both in their native range and have also overtaken production of native species in many countries including, for example, the UK, France, Ireland, and New Zealand. In 2007, Pacific oysters grown in these non-native ranges encountered a new variant of an Ostreid herpesvirus, termed OsHV-1 µVar [2], to which they generally proved to be highly susceptible. OsHV-1 µVar has since caused, and continues to cause, devastating losses across infected regions, with over 90% loss of naive stock being common and over 50% losses in previously exposed stocks [3, 4].

OsHV-1 is a large double-stranded DNA virus, and a member of the Malacoherpesviridae family within the Herpesvirales order. Whilst distant to the other families within Herpesvirales, the infection process of OsHV-1 maintains many of the classical herpesvirus features including a lytic phase, characterised by rapid onset, infection and mortality; and a latent (or persistent) phase, characterised by long-term low-level undetected infection. The lytic phase can further be characterised by expression of genes classified as immediate-early, early and late [5], each of which play specific roles in infection and replication, such as replication of viral DNA or production of virions. In susceptible animals, OsHV-1 commonly expresses groups of genes such as inhibitors of apoptosis proteins (IAPs) throughout infection [6], which inhibit the host immune response and promote viral replication. In addition, this immune suppression can lead to microbial dysbiosis and subsequent infection by opportunistic bacterial pathogens commonly present in the oysters’ natural environment. Together, these factors often result in mortality, the release of virions into seawater, and continuation of the infection cycle. Various studies have aimed to characterise the OsHV-1 infection process in oysters, e.g. [6–10], and there is broad consensus that the virus enters via the gill and/or digestive tract, before infecting large numbers of haemocytes and other parts of the body, with suggestions of some tissue tropism. However, the precise cellular basis of infection remains unresolved. Whilst most animals from susceptible populations will succumb to mortality in controlled conditions within around 96 hours, there is a notable amount of variation in resistance to infection. Some of this resistance is heritable, and there has been huge progress in breeding oysters with increased resistance to OsHV-1 through a combination of classical family-based selection and mass-selection. For example, mass selection in France [11], family selection or genomic selection in New Zealand [12], and family selection in Australia [13]. Resulting stocks have seen consistent increases in survival, including the extreme scenarios where mortality was reduced from 93% to 31% [14]. In addition, molecular approaches to breeding using a single quantitative SNP marker in the Pacific Northwest of the USA have resulted in lines with increased survival following exposure to local strains of OsHV1µVar [15]. However, given favourable conditions, the virus is still able to cause considerable losses in areas to which it has spread, and as such farmers have to take proactive measures to overcome losses, such as increased stocking at spat stages, or movement of animals away from OsHV-1 hotspots during periods of heavy infection.

Single-cell RNA-seq technologies are transforming bivalve and mollusc research, enabling high-resolution mapping of cell-type diversity [16–19], developmental trajectories [20, 21], germline specification [22], stress responses [23], and specialized functions such as biomineralization [24] and pigmentation [25], previously uncharacterised or underappreciated in bulk RNA-seq studies. In Pacific oyster, existing single-cell RNA-seq studies are limited to immune cells or select tissues, with major tissues such as gill and mantle, which constitute the bulk of oyster biomass and serve as the first point of contact with pathogens, not yet characterised at single-cell resolution. A comprehensive atlas representing major tissues and organs in Pacific oysters is lacking to-date. Such an atlas would provide a foundational reference for identifying cell types and their transcriptional programs across major tissues in Pacific oysters and other molluscs. It would also enable deeper insights into tissue-specific immune responses, development, and environmental adaptation, supporting a wide range of functional and comparative studies.

In this study, we aimed to generate a single-nucleus transcriptomic atlas of juvenile Pacific oysters during OsHV-1 infection. We applied single-nucleus RNA sequencing (snRNA-seq) to whole juvenile oysters which had been exposed to viral stock for up to 96 hours using a bath-challenge system. We sought to construct a high-resolution atlas of major cell types, resolve cell type-specific responses to viral exposure over time, identify populations potentially involved in viral entry, and determine the main cellular sites of viral replication through viral transcript detection. The resulting atlas provides high-quality single-nucleus markers for 17 of the 18 transcriptomic clusters resolved in our dataset, spanning major tissues and circulating immune cells. Together, these analyses establish a robust cellular framework for molluscan biology and provide new insight into host-pathogen interactions at single-nucleus resolution.

## Results

### snRNA-seq analysis reveals 18 transcriptomic clusters in juvenile Pacific oysters

To generate a snRNA-seq atlas of Pacific oysters during OsHV-1 infection, juvenile oysters were bath-challenged with the OsHV-1 µvar isolate [26] (Fig. 1A); uninoculated oysters exposed to seawater or uninfected oyster homogenate served as controls. Daily monitoring of mortality showed that 50% of oysters died within 96 hours of infection (Fig. 1B, top panel). Surviving oysters were sampled at 6, 24, 72, and 96 hpi for snRNA-seq library preparation. Viral infection was first evaluated in frozen tissue sections by qPCR (Fig. 1B, lower panel). Several samples exhibited relatively low levels of viral DNA, reflecting natural variability in the bath-challenge model. OCT-fixed whole-oyster samples with higher viral loads (see Fig. 1C) were used to generate snRNA-seq libraries, which were sequenced on an Illumina NovaSeq X Plus system.

**Figure 1.**
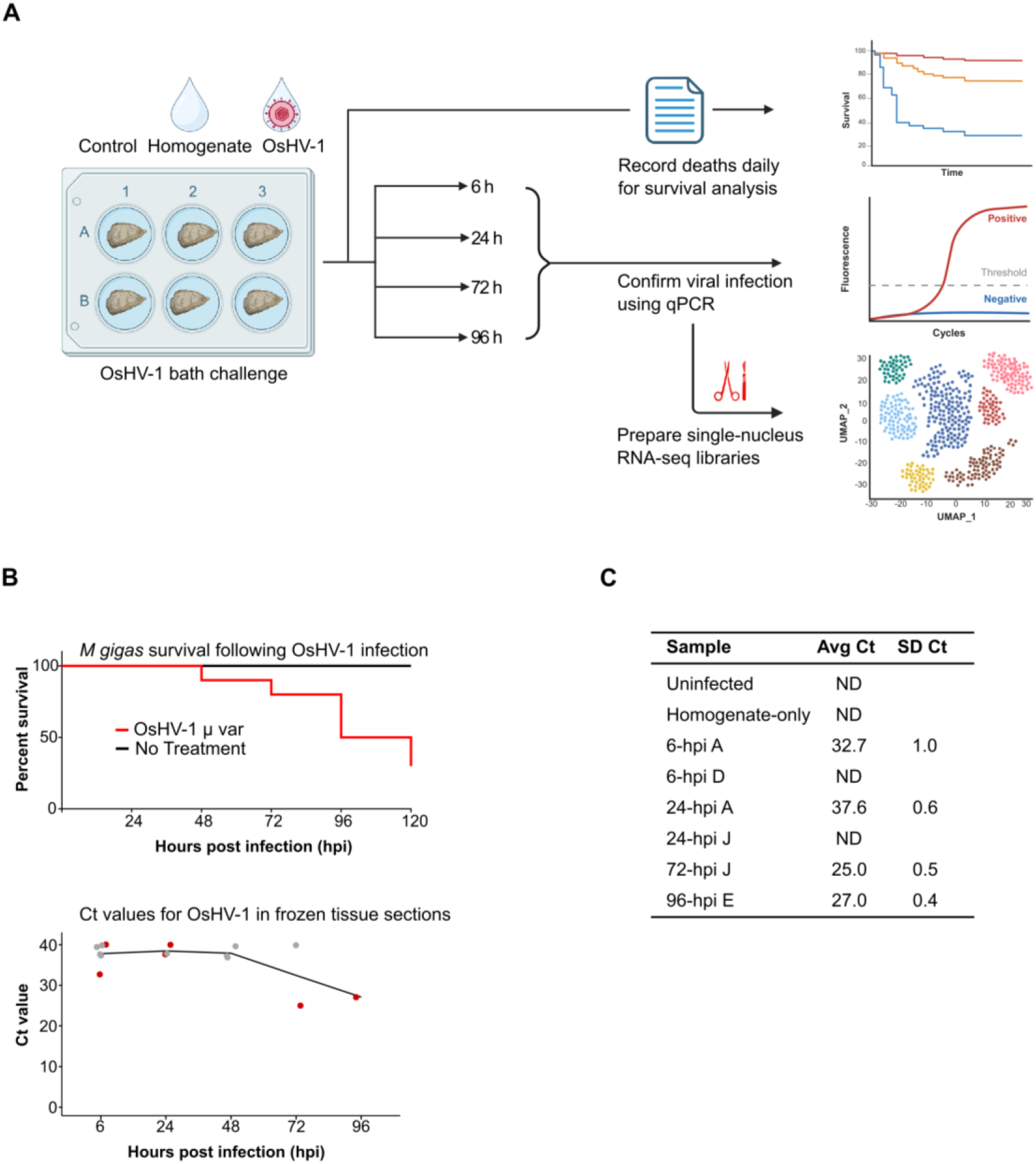
Schematic of the OsHV-1 Pacific oyster infection model, survival analysis, and viral load assessment for single-nucleus RNA-sequencing (snRNA-seq). **(A)** Schematic of the OsHV-1 bath challenge experiment showing the time points, in hours, at which oysters were sampled for survival analysis, qPCR-based detection of viral infection, and snRNA-seq library preparation. Infected oysters, as confirmed by qPCR assay, were used for the library preparation. **(B)** Upper panel: Survival curves for uninfected and OsHV1-infected oysters over 120 hours (five days). Lower panel: OsHV-1 DNA abundance in oyster samples across the infection time course. Individual dots represent qPCR Ct values for oyster samples collected at each time point. Samples selected for snRNA-seq library preparation are highlighted in red. The solid line indicates the mean Ct value at each time point; lower Ct values reflect higher viral DNA abundance. **(C)** Sample identifiers for the oysters that were selected for snRNA-seq library preparation, along with average Ct values and associated standard deviations for OsHV-1 µvar detection.

A total of 3.2 billion reads were generated across eight samples with 92% valid barcodes, yielding a sequencing saturation of 91% (Additional File 1 S1). An overview of the snRNA-seq bioinformatics pipeline is provided in Additional File 2 Fig. S1. After quality control filtering, a total of 21,189 nuclei were retained across eight samples, with a mean of 3,607 transcripts and 790 genes detected per nucleus (Additional File 2 Fig. S2). Integration of all samples enabled the identification of 18 distinct transcriptomic clusters consistently detected across all time points (Fig. 2).

**Figure 2:**
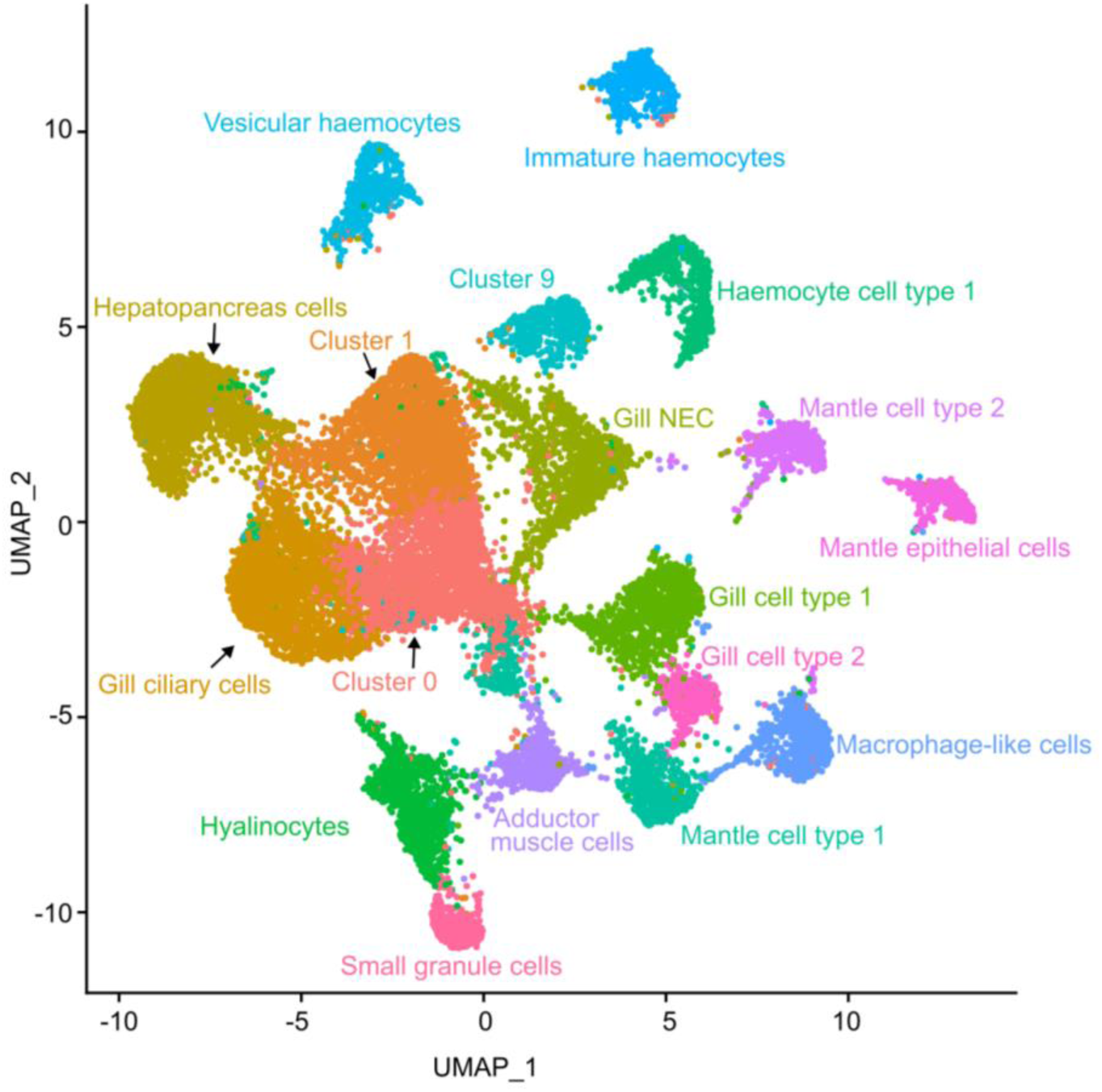
UMAP visualisation of a single-nucleus transcriptomic atlas of the Pacific oyster following OsHV-1 infection. Uniform Manifold Approximation and Projection (UMAP) visualization of ∼21,000 nuclei sorted into 18 transcriptomic clusters across eight samples. Uninfected and homogenate-only oysters served as experimental controls, while OsHV-1–infected oysters from a batch-challenge model were sampled at 6, 24, 72, and 96 hours post-infection.

### Supervised cluster annotation resolves haemocytes, gill cells, and mantle cells as dominant cell types in the snRNA-seq dataset

To assign cell-type identities to the 18 transcriptomic clusters (Fig. 2), we leveraged two publicly available Pacific oyster genome annotations. [19, 21]. By combining these two annotations, we were able to find either a gene symbol or protein domain description for 95% entries in our marker gene list (511/540 marker genes for the 18 transcriptomic clusters). Based on published cell-type marker reports from various bulk and single-cell RNA-seq studies in bivalve and fish species, we were able to characterise 15 of the 18 transcriptomic clusters. These included sub-populations of immune cells (haemocytes), gills, mantle, adductor muscle; three ambiguous clusters (cluster 0, 1, and 9) could not be assigned to any known cell type. The top two markers for all clusters are shown in Fig. 3 and Additional File 1 S2. Key marker genes used for cell type assignment are described below and shown in Additional File 2 Fig. S3, and the top 30 markers for each cluster are provided in Additional File 1 S3. Cell types are not always precisely identifiable, as such several classes are named by parent tissue-type (e.g. Mantle cell type 1), as identifiable from published tissue specific bulk RNA-seq.

**Figure 3.**
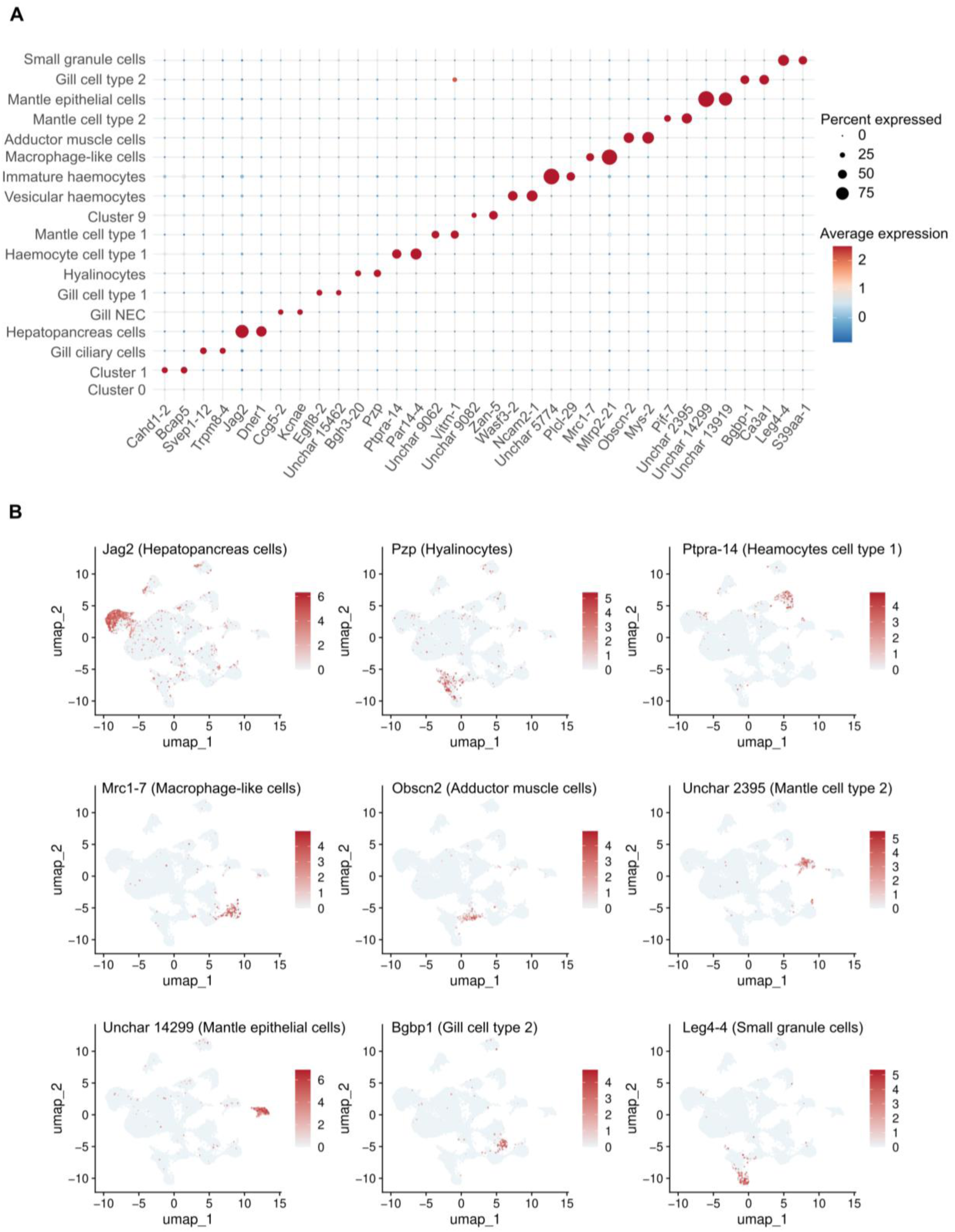
Top marker genes for the 17 transcriptomic clusters. For each cluster, the two most specific differentially expressed genes were selected as markers, defined by higher expression in the target cluster relative to all others (pct.1 = 0.24–0.96; pct.2 = 0.00–0.07; pct.1/pct.2 = 7–230). Cluster 0 did not yield any specific marker genes and was, therefore, excluded from the plots. (A) Dot plot: Dot colour indicates the average expression level of each gene within a cluster, while dot size represents the percentage of nuclei expressing the gene. (B) Feature plot: Expression patterns of representative marker genes for selected clusters visualised on a UMAP dimensionality reduction plot.

We assigned four transcriptomic clusters as gill cells in our datasets (Additional File 2 Fig. S3A). Gill ciliary cells (GCCs) were identified based on the expression of Kif19-a kinesin family protein localized to cilia tips [27], Trpv6-a calcium transporter enriched in gill neuroepithelial cells [28], and Trpm8- a cold and menthol-sensitive receptor expressed in ∼30% of cells at ∼21-fold higher levels than in other clusters. In addition, monocarboxylate transporter 14 (Slc16a), which has previously been identified as gill marker in zebrafish [29], was highly expressed in 56% of cells within this cluster. A distinct population of gill neuroepithelial cells (GNCs) was also detected based on the expression of voltage-gated potassium and calcium transporter family genes, together with other previously reported gill neuroepithelial markers [29]. Within this cluster, four potassium channel family genes (Kcnae, Kcnab, Kcnal-2, and Kcnaw-10) showed strong upregulation (log₂FC > 3.88). The most highly expressed gene was the voltage-dependent calcium channel gamma-7 subunit (Ccg5), detected in 25% of cells at levels approximately 100-fold higher than in other clusters. In addition, five polycystin genes (Pk1l2-3, Pk1l2-7, Pkdre-4, Pkd1-5, and Pkd2-4), which are implicated in calcium homeostasis, were expressed at very high levels. Consistent with the sensory function of neuroepithelial cells, multiple neurosensory genes, including Pclo [30], Dscam [31], and Fstl5 [32], were also abundantly expressed in this cluster. Another cluster exhibited a heterogeneous set of gill cell markers. Mucin genes (Muc5AC-like X2 and Muc5B) were highly upregulated (log₂FC > 4.5) in 62% and 71% of cells, respectively, consistent with mucous cell identity [29]. Neuroepithelial markers (Slc6a5, Meis2) showed moderate-to-high expression; Ece-2, an endothelial marker, was expressed in ∼23% of cells. The top cluster markers were Egfl8 and Notc2. Rfx4/6 transcription factor, which has been implicated in ciliated sensory neuron differentiation in Drosophila [33], was also enriched in this cluster. Given this mixed signature, we designated this cell population as gill cell type 1 (GCT1). A distinct transcriptional population was enriched for the gill neuroepithelial marker Slc6a5 [29] and three cathepsin L genes Catl 1, Catl 4, Catl5 that were previously reported to be expressed in mussel gills [34]. Top markers included Clca3a1 (calcium-activated chloride channel regulator; 54% of cells, ∼170-fold higher than other clusters), Bgbp-1 (β-1,3-glucan recognition protein), and chitin synthase. Immune genes CD109 (73% of cells) and Crr1-like were also expressed. Two angiotensin-converting enzymes showed high expression (logFC 5.5) in >50% cells. Based on the expression of mixed gill signature genes, this subset was annotated as gill cell type 2 (GCT2).

Three clusters were annotated as mantle tissue-derived populations. The first population, annotated as mantle cell type 1 (MCT1), exhibited expression of several previously described shell matrix proteins [35], including a putative chitinase 3 (Chi 10), chitotriosidase-1 (Chia-4), a chitin-binding protein peritrophin, two collagen alpha chains, cadherin-23, and protocadherin Fat 4 isoform X3 (Additional File 2 Fig. S3A). Notably, this population also showed high expression of a GATA transcription factor and a Hedgehog protein, suggesting a potential role for Hedgehog signalling in shell matrix formation [36]. A second mantle-associated population was annotated as mantle cell type 2 (MCT2) based on the elevated expression of mantle tissue–specific genes. Pif-7, a shell matrix protein [37], was expressed ∼390-fold higher in 35% of cells, while mantle protein 10, a shell gland marker [21], was detected at high levels in one-third of cells. Notably, three Notch and Notch-like proteins ranked among the top five markers, showing ∼600-fold higher expression in 37–40% of cells. We also observed high expression of two chitin-binding type-4 domain proteins and a chitin synthase protein. A third population corresponded to mantle epithelial cells (MECs), exhibiting high expression of four shell matrix proteins-putative insoluble matrix shell protein 5-like, Epdr1, Chia4, and teneurin-2 [35]. Seven collagen genes and WISP1 were also upregulated, consistent with roles in shell matrix formation. In addition, two C-type lectin domain–containing genes (Plcl-6 and Pgca-15) were enriched, supporting their involvement in biomineralization.

Adductor muscle cells (AMCs) were characterised by high expression of canonical adductor muscle markers, including unc-89 [17, 38] (obscurin; expressed in ∼60% of cells with ∼250-fold enrichment), myosin heavy chain and myosin tail 1 domain–containing proteins [17], the adductor catch muscle protein Cnn3 [21], the myocyte marker troponin, as well as multiple titin and collagen and fibrillar collagen NC1 domain–containing proteins (Additional File 2 Fig. S3B). Hepatopancreas cells (HCs, Additional File 2 Fig. S3B) were resolved based on the expression of multiple markers included in a previously defined hepatopancreas-enriched gene set [39]. This population exhibited elevated expression of salivary glue protein Sgs-4, death domain–containing protein, tripartite motif–containing protein 2, suppressor of tumorigenicity 14 protein homolog, Slit 3 protein, and three genes encoding fibrocystin-L. Notably, the Notch ligands Jag2 and Dner were expressed in 77% and 60% of cells within this cluster, respectively. Jagged proteins have previously been implicated as critical regulators of hepatopancreatic duct lineage specification in zebrafish [40].

Three clusters could not be confidently assigned to any known cell identity. Cluster 1 exhibited a strong immune-related transcriptional signature, characterised by the expression of several metalloendopeptidases including Mlrp1, Mlrp2, Uvs2, two PI3K adapter proteins (Bcap4, Bcap5), a macrophage migration inhibitory factor (Mif-1), and several RING-type domain-containing proteins (Diap2, Birc3, Birc7). In addition to immune markers, this cluster also expressed genes typically associated with mantle and shell matrix formation, including two VWFA domain-containing proteins, two chitin synthase proteins, a von Willebrand factor D and EGF domain-containing protein, and hemicentin-1. Due to this mixed gene expression profile, we were unable to assign Cluster 1 to a specific cell type (Additional File 2 Fig. S3B). Cluster 9 exhibited high expression of two genes encoding the male gonad marker zonadhesin, the acrosomal exocytosis protein Rims2, and multiple calcium and potassium voltage-gated channels, suggesting a potential male gonad–associated cell type. However, none of the well-established germline cell markers [22] (Ddx4/Vasa, Piwil1, TDRD1, and Boll/Dazl) were detected. As the top three genes in this cluster remain uncharacterised, we were unable to assign a specific cell identity to cluster 9. Cluster 0 lacked a clear set of marker genes (Additional File 1 S3), suggesting it may represent a heterogeneous or uncharacterised cell population.

### Six distinct haemocyte sub-populations resolved in the snRNA-seq dataset

Recently, a study [19] identified single-cell transcriptomic markers for seven haemocyte sub-populations in Pacific oyster. We examined the expression of these markers across transcriptomic clusters and identified multiple haemocyte subpopulations exhibiting enriched and cell type–specific expression. (Fig. 4 and Additional File 2 Fig. S3C). Hyalinocytes (HYL) were identified based on elevated expression of transforming growth factor-beta-induced protein ig-h3 (Bgh3-20, 118-fold higher expression, 31% cells in the cluster), and murinoglobulin-2. Another population expressed markers corresponding to unassigned haemocyte sub-population “cluster 5” described in the study [19]. Tll2, a zinc-dependent metalloendopeptidase, was expressed at high levels in 53% of cells, alongside protein-tyrosine phosphatase (Ptpra-14) and an uncharacterised gene (Unchar-5724). Based on this expression profile, we designated this population as haemocyte type 1 (HT1). Immature haemocytes (IMH) were resolved based on strong expression of genes encoding C- and H-type lectin domain-containing pattern-recognition receptors. Three of the top five expressed genes were C-type lectins. Additional immune-related genes in this cluster included bactericidal permeability-increasing protein/lipopolysaccharide-binding protein, lysosome-associated membrane glycoprotein 1, and two H-type lectins. Consistent with the immature haemocyte gene signature [19], this subset also displayed high expression of multiple ribosomal proteins (40S and 60S subunits). Small granule cell (SGC) were characterised by highly specific expression of several well-established markers [19], including galectin (Leg4-4), zinc transporters (Zip10, Znt2), PNPLA domain-containing protein (Yl446-8), TGc domain-containing protein (Tgm1-1), and alpha-1,4-glucan phosphorylase (Pygm). Another population expressed two vesicular cell markers [19]- Mylk-4 and Unchar-11539. We also detected strong expression of Wiskott–Aldrich syndrome protein family member 3 (Wasf3), a myeloperoxidase (Plsp-1), an alpha-2-macroglobulin family protein, and multiple integrin beta genes and Ig-like domain-containing proteins, further supporting its classification as vesicular haemocytes (VH). Finally, we identified a macrophage-like phagocytic haemocyte sub-population, which showed high expression of macrophage mannose receptor 1 (Mrc1) [41], proactivator polypeptide (Sap-1), and glutathione S-transferase sigma class protein (Gst8-1), genes associated with phagocytic and lysosomal activity. The expression of additional immune and metabolic regulators such as Kruppel-like factor 15 (Klf15) and Dmbt1 further supports its classification as a macrophage-like cells (MLC). No conserved marker for the haemocyte group, expressed exclusively in these six haemocyte subpopulations, was found (Additional File 2 Fig. S4).

**Figure 4.**
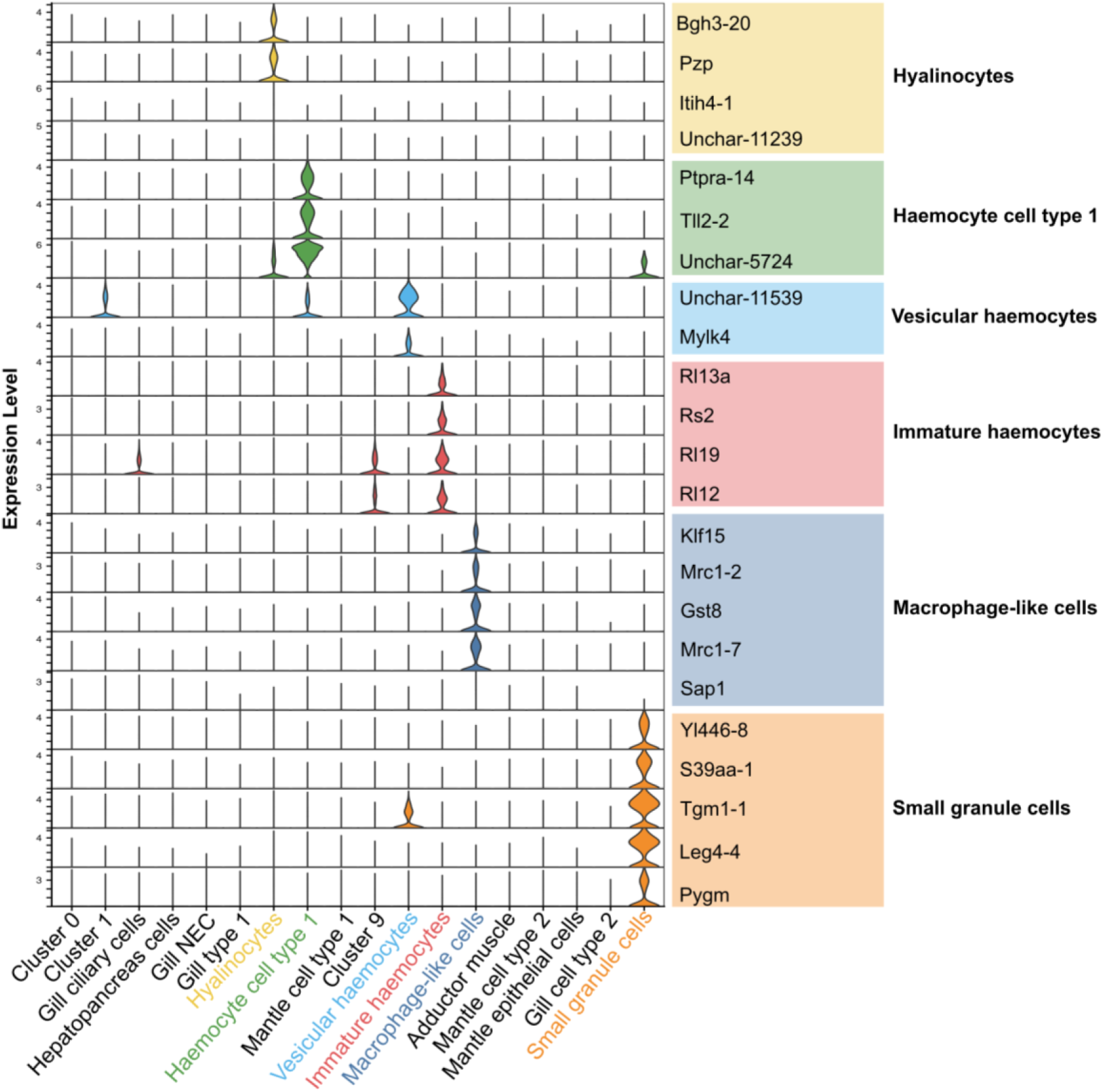
Haemocyte heterogeneity revealed by single-nucleus RNA-sequencing. We examined the expression of seven haemocyte sub-population markers reported previously (Divonne et al., 2025). Top 30 marker genes for each of these seven sub-populations were selected for expression across all 18 transcriptomic clusters in our dataset. Five haemocyte sub-types that were identified using Divonne et al datasets are: hyalinocytes, haemocyte cell type 1, vesicular haemocytes, immature haemocytes, and small granule cells. Markers for macrophage-like cells were selected based on previously reported literature, as described in the Results section. Only markers with high expression and strong sub-population specificity are shown. Violin plots on the left depict expression levels, panels on the right show key markers alongside the corresponding annotated haemocyte sub-types.

### OsHV-1 transcripts are detected nearly exclusively at 72 hpi

To monitor temporal expression of OsHV-1 transcripts during infection, we profiled viral gene expression across five conditions: uninfected controls and the four infection time points-6, 24, 72, and 96 hours post-infection (hpi). The OsHV-1 transcriptome, comprising 124 annotated open reading frames (ORFs), showed negligible expression in 6 and 24 hpi samples (Fig. 5). In contrast, high expression of viral transcripts was observed predominantly at 72 hpi (Additional File 2 Fig. S5), and limited expression in 96 hpi, consistent with viral load measurements obtained via quantitative PCR (Fig. 1B). At 72 hpi, viral transcripts were detected in multiple host cell types, including hyalinocytes, immature haemocytes, and notably, adductor muscle cells (Additional File 2 Fig. S5). The highest proportion of infected cells was found in the adductor muscle cluster, where approximately 9% of cells expressed OsHV-1 transcripts, followed by 8.1% in immature haemocytes and 6.6% in hyalinocytes (Additional File 2 Table S1). These findings suggest a broader cell tropism for OsHV-1 than previously recognised.

**Figure 5.**
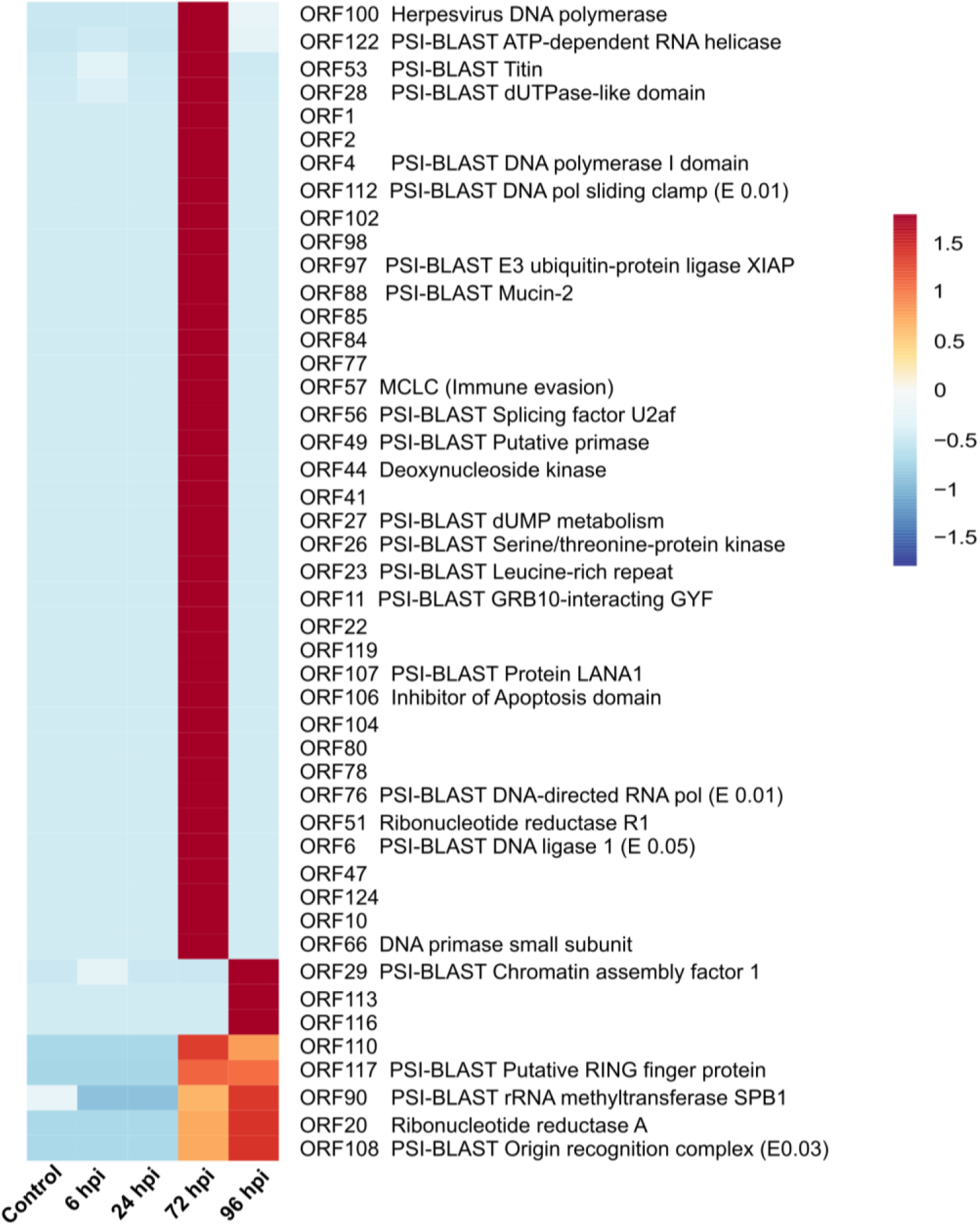
OsHV-1 transcripts are expressed in the 72 and 96 hpi samples. Expression of OsHV-1 ORFs across different infection stages are shown in the heatmap. Rows represent viral ORFs, columns represent oysters sampled at different time points after the OsHV-1 infection. Colour scale represents average gene expression per infection stage.

We utilized the OsHV-1 gene annotation provided by [5], and deployed PSI-BLAST to characterise previously unannotated ORFs. ORF100, encoding a DNA polymerase subunit, was expressed in all three major infected cell types—adductor muscle cells, hyalinocytes, and immature haemocytes—implicating these cells as primary sites of viral replication (Additional File 2 Fig. S5). In addition to ORF100, we detected expression of several ORFs involved in viral DNA replication, including ORF4 (DNA polymerase I domain protein), ORF49 (putative primase), ORF6 (DNA primase small subunit), ORF28 (dUTPase-like protein), ORF27 (involved in dUTP metabolism), and ORF51 (ribonucleotide reductase) (Additional File 2 Fig. S5). We also detected expression of immune modulatory genes-ORF57 (Mid-1-related chloride channel, MCLC) and ORF106 (inhibitor of apoptosis)-suggesting OsHV-1 may manipulate host immune responses to facilitate replication. Notably, these replication and immune modulation-associated transcripts were exclusively expressed at 72 hpi. Two DNA replication-related genes, ORF20 (ribonucleotide reductase) and ORF108 (origin recognition complex subunit), were expressed in both 72 and 96 hpi samples.

Capsid and packaging-associated transcripts showed a similarly restricted pattern, with ORF104 (major capsid protein) and ORF28 (portal/packaging-associated protein) detected predominantly at 72 hpi, especially in adductor muscle cells, consistent with active viral replication and progression to a productive stage of infection in this cluster (Additional File 2 Fig. S5). Expression was weaker in other infected populations, indicating that structural viral programmes were not uniformly distributed across cell types. Notably, ORF113 remained detectable at 96 hpi; given long-read evidence that it is co-transcribed with ORF112 [42], which encodes the VP19 capsid protein, this may indicate persistence of a structural or packaging-associated viral programme at later stages of infection.

### Differential gene expression (DGE) identifies infection response genes in Cluster 1

To understand the dynamics of cluster-specific transcriptional responses during OsHV-1 infection, we performed pairwise differential gene expression analyses within each cluster, contrasting control oysters with those at 6, 24, 72, and 96 hours post-infection samples. Three clusters showed differential expression of significant number of genes: cluster 1, hepatopancreas cells, and gill ciliary cells (Table 1, Fig. 6). Notably, cluster 1 had the largest number of differentially expressed genes (397 genes) in infected versus control samples (Additional File 2 Fig. S6). At 6 hpi, cluster 1 exhibited a pronounced innate immune activation, characterised by the upregulation of genes involved in inflammatory signalling and the initiation of programmed cell death (Fig. 7A, Additional File 2 Fig. S7). Elevated expression of Malt1, a CARD domain–containing protein, and the transcription factors Jun and Crebl2 indicates activation of the NF-κB and AP-1 pathways, key regulators of early antiviral and inflammatory responses (Additional File 1 S4). The concurrent induction of Fer, Abl2, Epha2, and Fgfr3 suggests early engagement of tyrosine kinase–mediated signalling, consistent with pathogen recognition and cytoskeletal reorganization in responding haemocytes [43]. Notably, the upregulation of Ced-3, a core executioner caspase, points to the onset of apoptosis in infected cells. In addition, the enrichment of multiple adhesion- and extracellular matrix (ECM)–related genes, including those encoding VWFA, fibronectin type III, and Ig-like domains, supports enhanced haemocyte aggregation and tissue remodelling associated with immune activation at this early stage.

**Figure 6:**
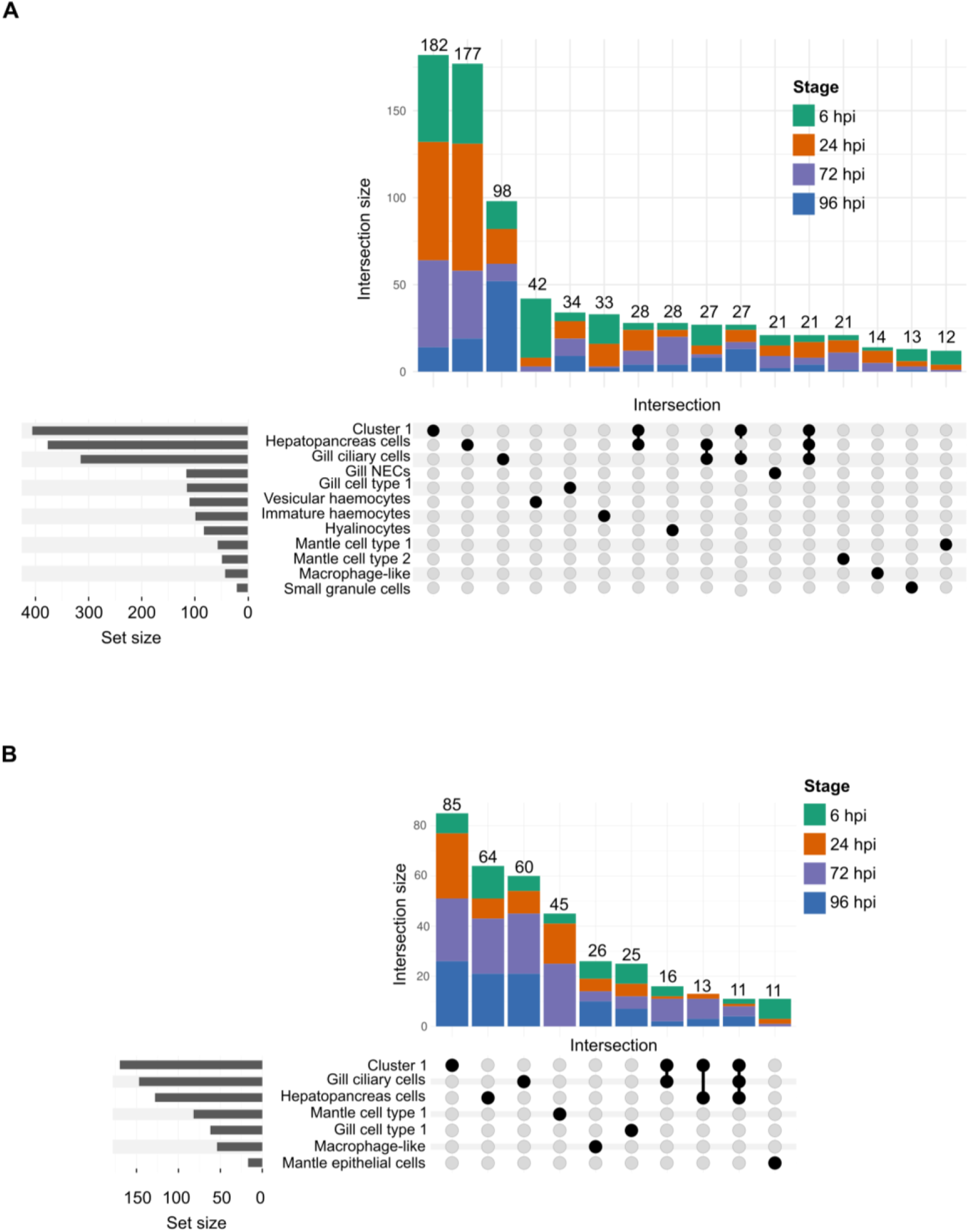
Intersection of differentially expressed genes among clusters across infection stages. UpSet plots showing overlaps of differentially expressed genes among transcriptomic clusters across infection stages. Genes upregulated **(A)**, and downregulated **(B)**, in the infected sample as identified by pairwise comparisons between control and infected samples at 6-, 24-, 72-, and 96-hours post-infection (hpi). Vertical bars indicate the size of gene intersections shared among clusters, with colours representing infection stages; only intersections with more than 10 genes are displayed. Horizontal bars on the left show the total number of differentially expressed genes per cluster. Black dots below the bars indicate clusters contributing to each intersection.

**Figure 7.**
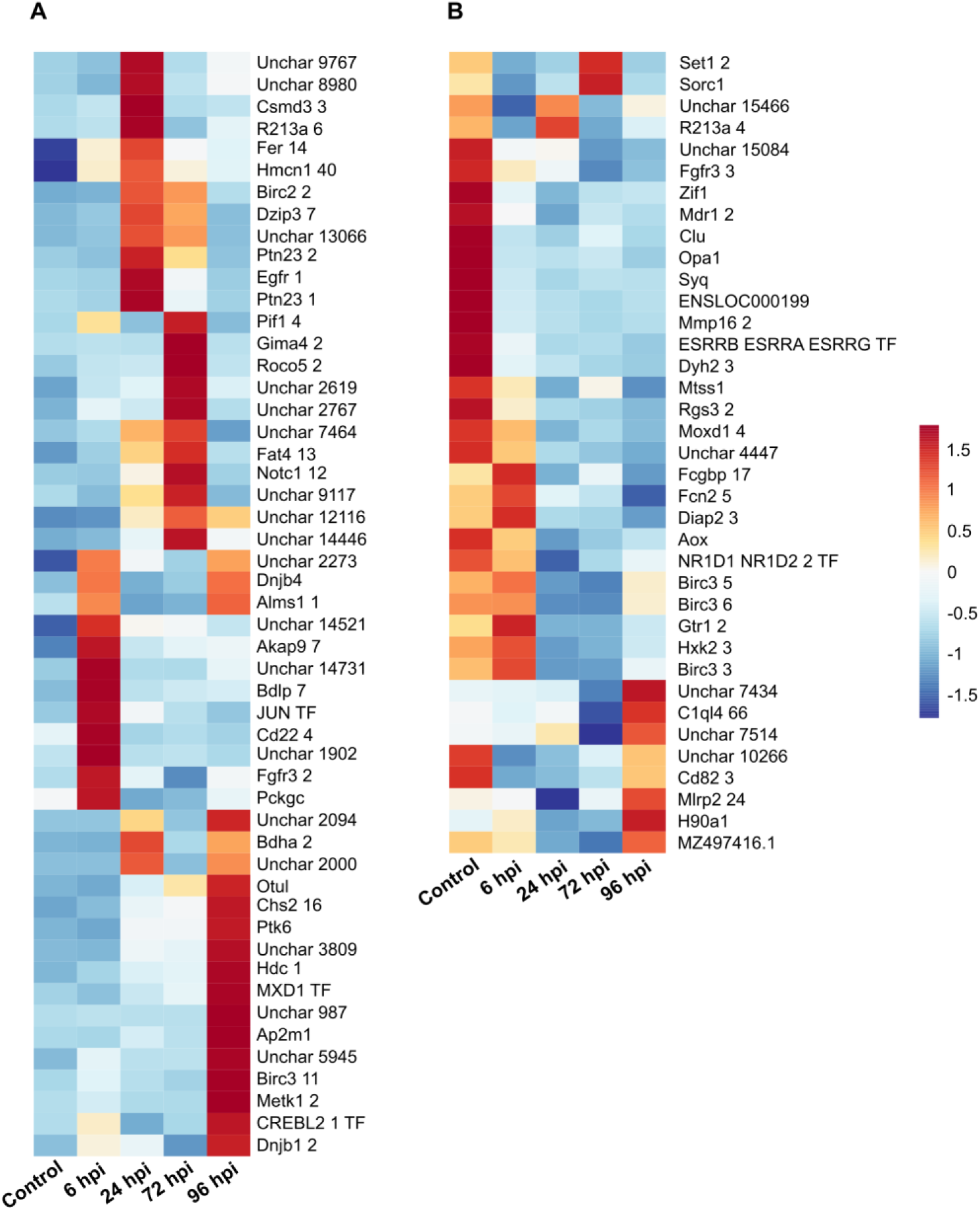
Differential gene expression (DGE) analysis identifies stage-specific immune modulation and apoptotic pathway activation in Cluster 1. Heatmap showing the average expression of top 15 genes per comparison that were identified as differentially expressed in cluster 1 across the four pairwise comparisons (control vs 6 hpi, control vs 24 hpi, control vs 72 hpi, and control vs 96 hpi). Expression patterns are shown across all five conditions to highlight temporal dynamics. Rows represent Pacific oyster genes, while columns represent samples. Genes upregulated **(A)**, and downregulated **(B)** in infected samples are shown in the heatmaps.

**Table 1.** Summary of differentially expressed genes across infection stages. Pairwise differential gene expression (DGE) analyses were performed for each cluster, comparing control samples with each infection stage (6, 24, 72, and 96 hpi). Columns show the number of differentially expressed genes for each comparison. The final column reports the total number of differentially expressed genes that were unique within cluster.

| Cluster identity | Infection stage |  |  |  | Total unique DEGs |
| --- | --- | --- | --- | --- | --- |
|  | 6 hpi | 24 hpi | 72 hpi | 96 hpi |  |
| Cluster 0 | 13 | 22 | 34 | 35 | 78 |
| Cluster 1 | 116 | 187 | 158 | 115 | 397 |
| Gill ciliary cells | 91 | 95 | 102 | 174 | 340 |
| Hepatopancreas cells | 117 | 149 | 127 | 112 | 382 |
| Gill neuroepithelial cells | 34 | 38 | 35 | 33 | 112 |
| Gill cell type 1 | 32 | 48 | 48 | 49 | 119 |
| Hyalinocytes | 16 | 19 | 26 | 26 | 70 |
| Haemocyte cell type 1 | 21 | 22 | 3 | 21 | 52 |
| Mantle cell type 1 | 29 | 53 | 44 | 13 | 104 |
| Cluster 9 | 15 | 16 | 8 | 29 | 53 |
| Vesicular haemocytes | 66 | 40 | 8 | 12 | 110 |
| Immature haemocytes | 34 | 42 | 12 | 29 | 95 |
| Macrophage-like cells | 15 | 36 | 21 | 25 | 68 |
| Adductor muscle cells | 6 | 9 | 6 | 20 | 33 |
| Mantle cell type 2 | 14 | 15 | 21 | 3 | 42 |
| Mantle epithelial cells | 16 | 9 | 6 | 2 | 29 |
| Gill cell type 2 | 7 | 25 | 12 | 10 | 40 |
| Small granule cells | 9 | 5 | 4 | 5 | 21 |

The 24 hpi stage of infection was characterised by a shift toward the induction of host factors involved in RNA sensing and degradation, indicating the activation of restrictive antiviral pathways. This was marked by the upregulation of cytosolic RNA sensors (Znfx1, Ddx58) and the concomitant induction of RNA-editing enzyme Dsrad 3 [44] and the exoribonuclease Helz2 (Additional File 2 Fig. S7), suggesting enhanced detection and degradation of viral dsRNA. Furthermore, we observed elevated expression of key signalling adaptors and regulators, including Traf3, TRIM family proteins, and multiple ubiquitination pathway enzymes (e.g., Ubp8, Ubp25, Dzip3, and E3 ligases), consistent with the robust activation of ubiquitin-mediated regulatory mechanisms in antiviral innate immunity. Strikingly, we observed high expression of Egfr1 (Fig.7A), consistent with previous evidence that EGFR signalling supports productive bovine herpesvirus 1 infection in cell culture [45]. As the infection progressed (72 and 96 hpi), cluster 1 exhibited strong enrichment of genes associated with apoptotic signalling and cell death regulation (Additional File 2 Fig. S7). Both pro- and anti-apoptotic proteins were highly expressed, including the apoptosis inhibitors Birc2 and Diap2, the key executioner caspase Ced-3, and Traf3, a signalling adaptor involved in apoptotic pathways. The sustained enrichment of multiple TRIM proteins and RING-type E3 ligases points to ubiquitin-dependent regulation of caspase activity, whereas the induction of MXD1 and CREBL2 suggests transcriptional integration of stress and survival pathways. Persistent expression of ECM genes (Hmcn1, fibrillin, cadherin and protocadherin) was also noted implying parallel tissue repair and restructuring accompanying apoptosis-driven turnover.

In cluster 1, OsHV-1 infection caused progressive suppression of genes associated with the normal transcriptional state. At 6 hpi (Fig. 7B), infected cells already showed lower expression of genes associated with mitochondrial organisation and function (Clu, Opa1), membrane transport (Mdr1, Zif1), DNA maintenance (Rev3l, Rbbp6), and transcriptional regulation, including HIF1A-like and ESRRG. By 24 hpi, this pattern had become more pronounced, with reduced expression of additional genes linked to metabolism and mitochondrial activity (Hxk2, Pckgc, Syq, Clu, Opa1), solute transport (Mdr1, Gtr1, Lat2), and regulatory pathways (Set1, CREBL2, NR1D-like, Sesn1). This reduction persisted at later time points. At 72–96 hpi, infected cluster 1 cells continued to show diminished expression of genes involved in mitochondrial respiration and organisation (Aox, Clu, Opa1), intermediary metabolism and transport (Hxk2, Pckgc, Syq, Mdr1), signalling (Ptprz, Rgs3), and dynein-associated structural functions. In addition, several genes with putative roles in cellular defence and stress responses, including Tbk1, NFAT5, Vdr, Fcn2/Fcgbp-like, and inhibitor-of-apoptosis family members (Birc3/Diap2-like), were also expressed at lower levels in infected oysters, particularly at 96 hpi. Overall, the cluster 1 response was characterised by broad reduction of mitochondrial, metabolic, transport, and regulatory transcripts following infection.

### OsHV-1 infection drives sustained antiviral signalling and metabolic reprogramming in the hepatopancreas cells

In the hepatopancreas population, OsHV-1 infection elicited a rapid induction of signalling, immune, and extracellular matrix–associated genes. At 6 hpi (Additional File 2 Fig. S8A, Additional File 1 S5), we observed increased expression of adhesion and matrix components containing Ig, EGF, VWFA, and fibronectin type III domains (e.g., Fgfr3, hemicentin paralogs, Vwde, Heg1), together with innate immune and signalling factors including Tbk1, Ptpra, and the apoptotic caspase Ced3. Early upregulation of tripartite motif proteins and ankyrin repeat–containing genes further pointed to activation of ubiquitin-dependent immune pathways. By 24 hpi, receptor tyrosine kinases and phosphatases (e.g., Egfr, Ptprt, Ptn23) and chromatin modifiers (Kmt5a and other SET domain proteins) were induced, alongside metabolic enzymes such as glutamine synthetase and arachidonate 15-lipoxygenase, indicating coordinated immune-metabolic reprogramming. At 72–96 hpi, sustained upregulation of E3 ubiquitin ligases (Ttc3, Birc2, Birc3, Bric7), helicases (Pif1), exonucleases, reverse transcriptase–like elements, and stress response genes (e.g., small heat shock proteins, H90a1) was evident. Induction of lipid metabolic regulators (Plin2, Fa2h), vesicular trafficking components (Ap2m1), and transcription factors (e.g., CREBL2, MXD1) at 96 hpi suggests prolonged antiviral defence coupled to extensive remodelling of metabolism and intracellular transport in this digestive-immune tissue.

In contrast to the activation of immune pathways, OsHV-1 infection elicited a pronounced and sustained transcriptional repression of genes governing core metabolic and structural functions in the hepatopancreas cells (Additional File 2 Fig. S8B, Additional File 1 S5). As early as 6 hpi, transcripts encoding mitochondrial alternative oxidase (Aox), axonemal dyneins (e.g., Dyh5), extracellular proteases and matrix proteins (e.g., Mmp16, VWFD family members), scavenger and VPS10-domain receptors (Sorc1), and the nuclear receptor ESRRG were significantly reduced. This initial repression broadened at 24–72 hpi to include key metabolic enzymes (e.g., Pckgc, Hxk2, Gfpt1), core cytoskeletal components (α- and β-tubulins, actin), and signalling regulators, alongside a decrease in the expression of immune lectins and fibrinogen/C1q domain–containing proteins. By 96 hpi, this pattern evolved into a broad and sustained suppression of structural, metabolic, and differentiation programs, evidenced by the persistent downregulation of dynein heavy chains, tubulins, actin, mitochondrial transporters, and developmental transcription factors such as Pax6. Collectively, these patterns indicate that OsHV-1 infection in the hepatopancreas redirects cellular resources away from mitochondrial respiration, cytoskeletal maintenance, and epithelial homeostasis.

### Temporal gene expression response to OsHV-1 in gill ciliary cells

In gill ciliary cells, OsHV-1 infection triggered a rapid and sustained induction of genes associated with pathogen sensing, signal transduction, and ubiquitin-mediated regulation. As early as 6 hpi, we observed increased expression of innate immune receptors and signalling components (Additional File 2 Fig. S9A, Additional File 1 S6), including the LPS detecting innate immune receptor Tlr4, the interferon regulator IRF2, and the NF-κB–associated scaffold Malt1, together with multiple tyrosine kinases and phosphatases (e.g., Fer, Ptprq) and apoptotic regulators such as Ced3 and Diap2. Concomitantly, numerous extracellular and adhesion-related proteins containing Ig, EGF, VWFD, and cadherin domains (e.g., Hmcn1, Ncam1, Vwde, Fat/Cadherin family members) were upregulated, indicating remodelling of epithelial interfaces. From 24–72 hpi, this response expanded to include antiviral and RNA-processing factors (e.g., Dsrad), DNA repair and helicases (Blm, Pif1, Neil1), tripartite motif proteins, and multiple E3 ubiquitin ligases, alongside heat shock proteins (Hsp70 family). By 96 hpi, elevated expression of stress response genes, lipid metabolic enzymes (Plin2, Fa2h), transporters, and additional ubiquitin pathway components persisted, consistent with sustained antiviral defence, proteostatic control, and metabolic reprogramming in infected ciliated epithelia.

Infection was also accompanied by broad repression of genes linked to mitochondrial function, cytoskeletal organization, vesicular trafficking, and cell proliferation (Additional File 2 Fig. S9B, Additional File 1 S6). At 6 hpi, transcripts encoding mitochondrial dynamics and respiration factors (e.g., Aox, Opa1, Clu) and the drug transporter Mdr1 were reduced, together with several RING-type proteins and the nuclear receptor ESRRG. Repression of metabolic enzymes (e.g., Hxk2, Pckgc) and ER chaperones (Bip) was evident by 24 hpi. At later timepoints (72–96 hpi), structural components of motile cilia and cytoskeleton, including α-and β-tubulins (Tba1c, Tbb) and axonemal dynein (Dyh2), were consistently downregulated, alongside extracellular matrix and adhesion molecules (e.g., Svep1, Fgfr3, Vwde family members). Additional repression of proliferation markers (Ki67) and stress-associated factors suggests a progressive shift away from growth and ciliary motility programs toward a state prioritizing antiviral defence over epithelial homeostasis.

## Discussion

### A high-resolution single-nucleus transcriptomic atlas of juvenile Pacific oysters

Our analysis of over 21,000 high-quality nuclei provides a high-resolution atlas of juvenile Pacific oysters, including during OsHV-1 infection. We have identified 18 distinct transcriptomic clusters spanning immune, epithelial, muscular, and digestive lineages, likely representing the most abundant cell types within the whole oyster samples sequenced. Existing reports are limited in scope and have focused on isolated biological contexts—larval development [21], germ cell specification [20], and immune cell diversity [19]—without yet providing a comprehensive, integrated multi-tissue single-cell atlas of the Pacific oyster. This atlas provides a significant advancement over bulk tissue transcriptomic analyses and presents a framework for understanding biology, such as antiviral response, at single-nucleus resolution in a mollusc species.

### Cellular diversity in key oyster tissues

The gill epithelium emerged as one of the most heterogeneous compartments, comprising ciliary cells, neuroepithelial cells, and two mixed epithelial subtypes, highlighting its functional complexity. These populations were enriched for voltage-gated ion channels, polycystins, and calcium-regulatory genes, consistent with a specialised sensory interface. Given that herpesviruses commonly hijack host calcium signalling to promote entry and replication [46], gill sub-populations may constitute critical sites of early host–virus interaction. Consistent with this, gill ciliary cells ranked among the clusters with the highest number of infection-induced differentially expressed genes. The co-expression of mucins and immune recognition molecules further indicates tight coupling of barrier function and innate immunity at the host–environment interface.

Mantle tissue resolved into three transcriptionally distinct populations enriched for well-characterised shell matrix proteins [35], chitin-associated enzymes, and collagen family members, underscoring functional specialisation within biomineralising epithelia. Elevated expression of a GATA transcription factor supports its proposed role in shell formation (Gang Liu et al., 2015). Notably, hedgehog genes were co-expressed in GATA-positive cells, raising the possibility of coordinated activity between these pathways during shell matrix production.

Hepatopancreas populations displayed a transcriptional profile consistent with integrated metabolic, structural, and immune functions. Expression of Notch ligands Jag2 and Dner across this population suggests involvement of developmental signalling pathways associated with epithelial maintenance and plasticity. Such signalling activity may reflect heightened epithelial plasticity or regenerative responses triggered by infection. Our data indicate that the hepatopancreas may be an early site of OsHV-1 engagement [47], as reflected by its extensive transcriptional remodelling during infection. Adductor muscle cells in our dataset were annotated based on the expression of canonical muscle markers and adductor catch muscle protein, however, we cannot fully exclude contributions from other muscle cell types, such as those from the heart or mantle. Nevertheless, given the dominance of the adductor muscle in oyster tissue, this population most likely represents adductor muscle cells.

Three clusters remained unresolved, as mixed marker expression precluded confident assignment to a defined cell type or tissue of origin. This ambiguity may reflect the substantial transcriptional overlap characteristic of bivalve tissues: comparative analyses in mussels have shown that up to 55% of transcripts may be shared across tissues [34], and in Pacific oyster, gill tissue shares numerous transcripts with circulating immune cells [48]. Such overlap is consistent with the broad distribution of innate immune and physiological programmes in bivalves. Additionally, limited gene annotation may restrict the identification of cell-type specific markers, contributing to unresolved cluster identities. Recent work [49] demonstrated that haemocytes can undergo dynamic state changes, including degranulation, shifting between granulocyte and agranulocyte phenotypes. Such plasticity would be expected to generate mixed transcriptional signatures, as cells transiently co-express markers associated with multiple states. In this context, unresolved clusters may reflect biologically meaningful intermediate or activated states.

### Immune cell heterogeneity in Pacific oyster

We identified six haemocyte subpopulations, four of which showed expression highly congruous with previously established markers [19]. Notably, we identified a macrophage-like population not previously resolved in oyster single-cell studies. In bivalves, such cells are best defined by phagocytic and endolysosomal programmes rather than strict vertebrate marker conservation, and transcriptomic studies of purified oyster phagocytes have shown enrichment of antibacterial defence and phagocytosis-associated pathways [50]. Consistent with this, the macrophage-like cluster expressed mannose receptor C-type 1 (Mrc1), scavenger receptor- and lectin-like genes, Dmbt1, Fcn2/C1q-like transcripts, and multiple genes involved in lysosomal and vesicular trafficking, including Lyst, Lamp1, and Eea1, supporting a role in pathogen recognition, uptake, and intracellular processing. Given recent single-cell evidence of extensive haemocyte heterogeneity, this population may represent a specialised macrophage-like haemocyte subtype not previously resolved in conventional classifications [51].

It is likely that unassigned ‘Cluster 1’, which exhibited a strong response to OsHV-1 infection, represents an uncharacterised haemocyte subtype. Although canonical haemocyte markers [19] were not detected, this cluster displayed a pronounced immune-associated transcriptional signature, including high expression of Mif-1, a key regulator of innate immunity in mammals [52], together with genes linked to immune signalling and activation, such as phosphoinositide 3-kinase adapter protein 1, RING-type domain-containing proteins, RhoU, C2 domain-containing protein, subtilase, and multiple metalloendopeptidases. These features are consistent with an activated immune-like state and suggest that cluster 1 may correspond to a previously uncharacterised haemocyte-like population.

### OsHV-1 infection response genes

Only one oyster was analysed per infection condition, limiting robust statistical assessment of differential gene expression. Nevertheless, we identified immune response genes displaying distinct pro- and anti-viral signatures [6], including those associated with RNA surveillance and RNA editing. De Lorgeril et al. showed that OsHV-1 susceptibility in Pacific oysters is marked by delayed antiviral activation, strong mobilisation of IAP-associated anti-apoptotic pathways, and immune dysfunction. Against this framework and given that our experiment followed a single cohort that was likely susceptible overall, the infection-responsive genes identified here likely capture variation within a broadly susceptible host-response state. Gill ciliary cells displayed the earliest and clearest antiviral response at 6 hpi, with induction of IRF2, SOCS2, TLR4, BIRC3, DIAP2, MALT1, TRAF3 and DDX58-like genes, but simultaneously showed reduced expression of genes associated with ciliary structure and epithelial integrity. Hepatopancreas cells also mounted an early antiviral program, including TBK1, IRF2, TLR4, MyD88, BIRC2/BIRC3, TRAF3, DDX58-like and ZNFX1, while losing expression of genes linked to metabolic, scavenging and epithelial homeostasis, such as Pckgc, Aox, C49a1-like, Abca5, Sorc1 and lectin/scavenger-receptor-like factors. By contrast, cluster 1 was dominated by a more persistent and delayed response spanning 24–96 hpi, marked by TRAF3, DDX58-like, SAMHD1-like, ZNFX1, BIRC2/BIRC3, CYLD and OTULIN, together with reduced expression of mitochondrial, transport and homeostatic genes. Collectively, these patterns indicate that OsHV-1 infection induces both activation of antiviral and apoptosis/survival pathways and suppression of tissue-specific functions across multiple cell types.

### Virus replication and packaging in adductor muscle cells

In an *in vitro* haemocyte OsHV-1 infection model, Dotto-Maurel et al. classified herpesvirus genes into immediate-early, early, and late groups [5], but showed that transcription is largely cumulative rather than strictly temporally segregated. The predominance of OsHV-1 transcripts at 72 hpi in our datasets likely reflects the asynchronous nature of infection in the whole-oyster bath-challenge model, in which multiple tissues and cell types are sampled simultaneously and may occupy different stages of viral progression. Unlike synchronised *in vitro* haemocyte infections, whole-animal infection is expected to involve delayed dissemination, cell type-specific permissiveness, and substantial spatial heterogeneity, making a clear temporal transcriptional cascade difficult to resolve. Consistent with this, the 72 hpi sample contained viral ORFs assigned to multiple kinetic classes, including immediate-early genes such as ORF27, ORF80, ORF104 and ORF122, and early genes such as ORF100, ORF106, ORF20, ORF51, ORF49 and ORF66.

Similarly, at the cellular level, OsHV-1 transcription did not segregate into discrete immediate-early, early, or late gene sets, but instead showed broad co-expression of multiple ORFs within the same cell populations. Highly infected clusters, particularly adductor muscle cells and two of the haemocytes sub-populations (immature haemocytes and hyalinocytes), simultaneously expressed replication-associated, metabolic, and immune modulatory OsHV-1 ORFs, indicating that transcripts from different functional stages of the viral programme are detected together. We, therefore, interpret the 72 hpi timepoint as a composite snapshot of productive infection, reflecting asynchronous viral progression across tissues.

Notably, adductor muscle, which has previously been suggested as a primary site of viral replication [7], showed the clearest signature of productive infection, with concurrent expression of replication-associated ORFs (ORF100, ORF4, ORF49, ORF51 and ORF27) together with capsid- and packaging-associated ORFs, including ORF104 and ORF28. This is especially striking given that infection was established by bath challenge in our study rather than direct injection into the adductor muscle, further supporting the adductor as a primary site of viral replication. Unexpectedly, only a small number of host genes were differentially expressed in adductor muscle during infection. Finally, ORF113 was enriched in mantle cell type 2 at 96 hpi; this is of particular interest given long-read evidence that ORF113 is expressed as part of a polycistronic transcript with ORF112 [53], which encodes the VP19 capsid protein, suggesting that structural viral transcription may persist in a more cell type-restricted manner.

### Challenges and limitations

Our datasets likely captured the most abundant cell types within each major tissue, whereas less abundant populations—or those less amenable to nuclei extraction- were underrepresented and not captured. This occurred despite high sequencing saturation (∼90%) and ∼3000 nuclei profiled per sample, suggesting that overwhelming number of cell types recovered from whole oysters may require the sampling of more nuclei per sample to be fully elucidated. Alternatively, future studies exploring individual tissues at single-cell level will result in a higher number of nuclei per cell type and will further identify more sub-types. For example, gill [18, 28, 29] and adductor muscle [17] tissues in bivalves may each comprise more than 20 distinct subpopulations, highlighting the likely underestimation of cellular diversity in our dataset.

As with other bivalve species, poor gene annotation poses a significant bottleneck in Pacific oyster research, many of the key markers identified in our dataset remain either annotated only with generic protein domains or completely uncharacterised. Although, most reads mapped to the reference genome (∼80%, of which 22% multi-mapped), only a smaller proportion could be uniquely assigned to annotated genes (∼43%). This likely reflects limitations of the current Pacific oyster genome resources, as the species has a highly polymorphic and repeat-rich genome [54], leading to ambiguous read mapping. Future efforts to improve functional annotation will not only refine gene models but also enable the broader scientific community to more accurately identify and characterise cell types, regulatory networks, and key OsHV-1 response pathways in Pacific oysters.

## Conclusions

Taken together, we present a foundational cellular framework for understanding host–virus interactions in juvenile oysters during OsHV-1 infection. Beyond refining oyster cell-type annotation, our findings suggest that antiviral response in bivalves is distributed across specialised tissue compartments rather than confined to haemocytes. Tissue-resolved spatial and viral transcriptomic analyses in the future will further uncover oyster cellular heterogeneity and delineate the precise entry portals and replication compartments exploited by the virus.

## Methods

### Experimental design and setup

Juvenile “G10” graded Pacific oysters (*Magallana gigas*), sourced directly from Guernsey Sea Farms (Helm, Guernsey), were acclimated in 28 ppt 0.22 µm-filtered artificial seawater (ASW) before being transferred to 22 6-well plates, with each well containing 7 mL of 28 ppt 0.22 µm-filtered artificial seawater (ASW). One oyster was placed in each well, and the plates were incubated at 20 °C.

### Preparation of OsHV-1 stock

OsHV-1 was obtained from a preserved master stock “MS8” (approximately 3×10^5^ viral copies/mL) from CEFAS laboratory (Weymouth, UK). This strain, originally isolated from an outbreak in Poole Harbour, Dorset, UK and passaged twice through laboratory animals, has also been genomically characterised [55]. To amplify the virus, the largest oysters (10-20 mm) were anaesthetized in seawater containing 50 g/L magnesium chloride for 4 hours. Oysters were then grouped and injected with 50 µL OsHV-1 MS8 stock or artificial seawater as a control. Mortality was checked daily, and water was changed every 48 hours. Dead or moribund oysters were stored at 4 °C. At the end of the challenge, gill and mantle tissues were removed from dead oysters, homogenised, diluted in ASW, and sequentially filtered through syringe filters (5 µm > 1 µm > 0.4 µm > 0.2 µm). The prepared virus (MS9) and an equivalent homogenate control without virus were mixed with 10% glycerol as a cryoprotectant and stored at −80 °C prior to use.

### Whole animal disease challenge

To prepare for the infection, MS9 OsHV-1 virus and homogenate control were thawed on ice. The virus solution was prepared by combining the contents of the OsHV-1 with 480 mL of 28 ppt ASW and approximately 40 mL of algae, resulting in a final amount of approximately 2×10^5^ viral copies per well. The homogenate solution was prepared by mixing an equivalent ratio of homogenate to 96 mL of 28 ppt ASW and approximately 4 mL of algae.

For the infection challenge, ASW in each well of 6-well plates was replaced with freshly prepared ASW, supplemented with either the prepared OsHV-1 virus solution or the homogenate control solution. There were ten replicates for each time point of viral challenge and two replicates at each time point for homogenate control. Two empty wells in one plate were filled with the OsHV-1 virus solution. Oysters were checked for mortality daily, and water was replaced every 48 hours.

Samples were collected at time zero (T0), and at 6, 24, 48, 72, and 96 hours post infection (hpi). These samples were frozen in optimal cutting temperature medium (OCT, VWR, UK) on a bed of dry ice and then stored at −80 °C. Water samples were collected from each well prior to shucking the oysters. Up to 24 hpi, both the infected and homogenate control oysters were sampled, with no mortalities observed at this stage. At 48 hpi, the oysters and homogenate controls were sampled again; some oysters were slow to close but no deaths were noted in these plates. At 72 hpi, mortalities were observed in four wells. At 96 hpi, mortalities had been recorded for seven wells, as well as in a single homogenate control well.

### Infection quantification

Approximately ten 20 µm thick sections of tissue were cut from each sample using a cryostat at −20 °C before returning samples to −80 °C storage. DNA was extracted from each batch of tissue sections with a Qiagen Blood and Tissue kit before normalisation and quantification with the TaqMan qPCR assay derived from [56]. Briefly, 2 µL sample template was mixed in a 20 µL reaction with 10 µL 2X NEB Luna master mix, 0.4 µM Forward primer OsHV-1 BF (5’ GTCGCATCTTTGGATTTAACAA 3’), 0.4 µM reverse primer OsHV-1 B4 (ACTGGGATCCGAACTGACAAC 3’), and 0.2 µM OsHV-1 probe (5’ FAM-TGCCCCTGTCATCTTGAGGTATAGACAA-TAM 3’) and then amplified in an Applied Biosystems 7500 qPCR system [8]. One sample from each of the seawater control and negative homogenate control groups was selected for single-nucleus RNA-seq, alongside six virus-exposed samples: two from 6 hpi, two from 24 hpi, and one each from 72 hpi and 96 hpi. Exposed samples had a mixture of high and low infection levels, aiming to capture variation in the timeline of infection (Fig. 1).

### Single-nucleus library preparation and sequencing

Nuclei were isolated from juvenile oysters using an adapted protocol originally optimised for Atlantic salmon epidermis [57]. Briefly, juvenile oysters were minced for 5 min, chilled on ice, and further homogenised by four successive rounds of scissor chopping in 1 mL TST buffer until a homogeneous suspension was obtained. The lysate was then filtered through a 40-µm Falcon cell strainer. Subsequently, 1 mL TST buffer and 3 mL 1× ST buffer were added to each sample, followed by centrifugation at 500 x g for 5 min at 4 °C. The pellet was resuspended in 1 mL of 1× ST buffer and passed through a 40-µm cell strainer twice more. Nuclei integrity was assessed by haemocytometry with Trypan Blue staining and by fluorescence microscopy. Nuclei concentration was then estimated using a disposable flow haemocytometer (C-Chip Neubauer Improved, 100 µm depth; NanoEnTek, catalogue no. DHC-N01).

Isolated nuclei were fixed with the Parse Biosciences Nuclei Fixation Kit V2 (Seattle, WA) according to the manufacturer’s protocol and stored at −80 °C. Single-nucleus RNA-seq libraries were prepared using the Parse Biosciences Evercode WT Standard Kit V2.1, targeting 7,500 nuclei per sample. cDNA was amplified for 12 PCR cycles, followed by 8 cycles of index PCR. Library concentration was measured using a Qubit v3 High Sensitivity DNA assay (Thermo Fisher, Waltham, MA), and library quality was assessed on a TapeStation 4200 with the High Sensitivity D5000 assay (Agilent, Santa Clara, CA). Sequencing was performed on an Illumina NovaSeq X Plus System with a 25B flow cell using paired-end 150 bp reads, aiming for approximately 400 million reads per library.

### Single-nucleus RNA-seq data analysis

#### Reference genome and annotation

A composite genome for the host and pathogen was created by concatenating Pacific oyster genome assembly cgigas_uk_roslin_v1 [54] and OsHV-1 virus genome [26]. Similarly, a composite GTF annotation file was created by combining Crassostrea_gigas_uk_roslin_v1.gtf and viv46-2-m_assembly_NR_final_ok.gtf files.

#### Data processing and quality control

Raw FASTQ files were evaluated with FastQC and summarised using MultiQC. Sequencing metrics were broadly consistent across libraries, with fixed read lengths (150 bp) and stable GC content. We obtained significantly lower rates of transcriptomic mapping with Parse Biosciences proprietary tool ‘splitpipe’, therefore, we used STARsolo tool version 2.7.10b [58] for read alignment. The barcode file from splitpipe output (combine output > process > barcode_data.csv) was, however, used to create whitelist files for mapping with STARsolo. FASTQ.gz files for each of the eight Parse sub-libraries were aligned to the composite genome using STARsolo with --outFilterScoreMinOverLread 0.50 and --outFilterMatchNminOverLread 0.50 parameters. On an average, 354 million reads per sample were used as input, of these 80% aligned to the genome; 55% of reads mapped to GeneFull (exons and introns); median reads and UMI per nucleus were 54,215 and 5300, respectively. STARsolo output files UniqueAndMult-EM.mtx, barcodes.tsv, and features.tsv in the GeneFull raw directory were used to create Seurat object for each sub-library using Seurat 5.0 [59]; Parse Biosciences Evercode chemistry uses random hexamers and poly(A) oligos to capture mRNA, these two were merged prior to making Seurat objects for the correct assignment of nuclei.

Single-nucleus RNA-seq data from the eight sub-libraries were processed in R (v4.2.2) using Seurat. Nuclei were first filtered within each Seurat object to remove likely low-quality nuclei and outliers, retaining only those with between 200 and 3,000 detected genes, and excluding nuclei with more than 5% mitochondrial UMIs. Ribosomal RNA genes were also removed from the expression matrices. The filtered Seurat objects from the eight sub-libraries were then merged prior to standard downstream analysis.

#### Normalisation, dimensionality reduction, and clustering

Samples were normalised using LogNormalize method with the default scale factor 10,000. Highly variable genes were identified using the vst method with nfeatures set to 3000, which were used for dimensionality reduction using Principal Component Analysis (PCA). Uniform Manifold Approximation and Projection analysis (UMAP, [60]) was performed to visualize the data with these settings: dims = 1:18, resolution = 0.6. The original Cluster 1 in our dataset had unusually high proportion of mitochondrial reads and a diffuse, non-specific marker gene expression profile. Therefore, we removed this cluster from the original seurat object using ‘subset’ function and datasets were re-analysed with the same parameters as mentioned above for all the steps from normalisation to UMAP analysis. A total 33,466 nuclei were identified with 21,189 genes detected across eight samples. Genes were annotated by combining comprehensive annotations from [19, 21]. To identify marker genes for clusters, we used Seurat’s FindAllMarkers function (min.pct = 0.20, log2fc.threshold = 0.25). For cell type assignment, top 30 expressed genes in each cluster were extracted using Extract_Top_Markers function in scCustomize [61] version 3.0.1.

#### Pairwise differential gene expression analysis

Cell-type identities were integrated with condition labels (control and 6-, 24-, 72-, 96-hpi groups) to define sample subtype–specific groups, enabling matched, cell-type–resolved analyses. Differential gene expression was performed using Seurat (FindMarkers), comparing control nuclei with nuclei from the corresponding infected samples at each time point. Genes expressed in at least 25% of nuclei in either group (min.pct = 0.25) were tested using a minimum log2 fold-change threshold of 1, and analyses were conducted independently for each control–time point comparison.

## Abbreviations

ASW: Artificial Seawater
DEG: Differentially Expressed Genes
DGE: Differential Gene Expression
HPI: Hours Post-Infection
IAP: Inhibitor of Apoptosis
ORF: Open Reading Frame
OsHV-1: Ostreid Herpesvirus 1
PCA: Principal Component Analysis
snRNA-seq: Single-Nucleus RNA Sequencing
UMAP: Uniform Manifold Approximation and Projection Analysis
UMI: Unique Molecular Identifiers
VWFA: Von Willebrand Factor Type A

## Additional files

**Additional File 1.xlsx**

This Excel file contains six sheets: **Sheet 1:** STARsolo alignment summary; **Sheet 2:** Top two markers for each transcriptomic cluster; **Sheet 3:** Top 30 marker genes for each transcriptomic cluster; **Sheet 4:** List of differentially expressed genes in ‘Cluster 1’ in response to infection; **Sheet 5:** List of differentially expressed genes in ‘Hepatopancreas Cells’ in response to infection; **Sheet 6:** List of differentially expressed genes in ‘Gill Ciliary Cells’ in response to infection.

**Additional File 2.docx**

This word document contains the following figures:

Figure S1. Bioinformatics workflow for single-nucleus RNA sequencing analysis.

Figure S2. Quality Control metrics across samples in the merged single-nucleus RNA-sequencing datasets.

Figure S3. Expression of key marker genes across transcriptomic clusters described in the Results section.

Figure S4. Conserved marker expression in the haemocyte group.

Figure S5. Expression of OsHV-1 transcripts across samples.

Figure S6. Heatmaps of differentially expressed genes in clusters with the highest transcriptional response to infection.

Figure S7. Immune modulatory genes upregulated in cluster 1 in response to OsHV-1 infection.

Figure S8: Top genes differentially expressed in the cluster hepatopancreas cell type.

Figure S9: Top differentially expressed genes identified in the gill ciliary cell cluster.

Table S1. Distribution of nuclei expressing viral transcripts across cell subtypes and infection conditions.

## Declarations

### Ethics approval and consent to participate

Not applicable.

### Consent for publication

Not applicable.

### Data and code availability

Raw single-nucleus RNA-sequencing data have been deposited in the NCBI Sequence Read Archive (SRA) under BioProject PRJNA1401775. Individual sample data are registered under the following BioSample and corresponding SRA run accessions: SAMN54570629 (SRR36788949), SAMN54570630 (SRR36788948), SAMN54570631 (SRR36788947), SAMN54570632 (SRR36788946), SAMN54570633 (SRR36788945), SAMN54570634 (SRR36788944), SAMN54570635 (SRR36788943), and SAMN54570636 (SRR36788942).

All code used for data analysis is available at the Roslin-Aquaculture GitHub repository (https://github.com/Roslin-Aquaculture/single-nucleus-RNA-seq-analysis-Mgigas).

### Authors’ contributions

TPB, TR conceived the idea, provided resources, and co-supervised the project; TR, TPB collected samples and set up disease challenge; AFC, JJF, AF, TR prepared snRNA-seq libraries; TR performed real-time PCR experiments; PSD led snRNA-seq data analysis with support from TCC, RST, TPB; PSD, TR, AFC, TPB co-produced figures and additional files; PSD, TPB wrote the first manuscript draft; all authors drafted, revised, and approved the final manuscript.

## Acknowledgements

The authors would like to thank Dr Aurélie Dotto Maurel for providing OsHV-1 genome and annotation data, and the wider Roslin institute aquaculture team for support during the process of producing this manuscript.

