## Additional File 2 for "A Single-Nucleus Transcriptomic Atlas Reveals Cell Type-Specific Responses to OsHV-1 Infection in the Pacific Oyster"

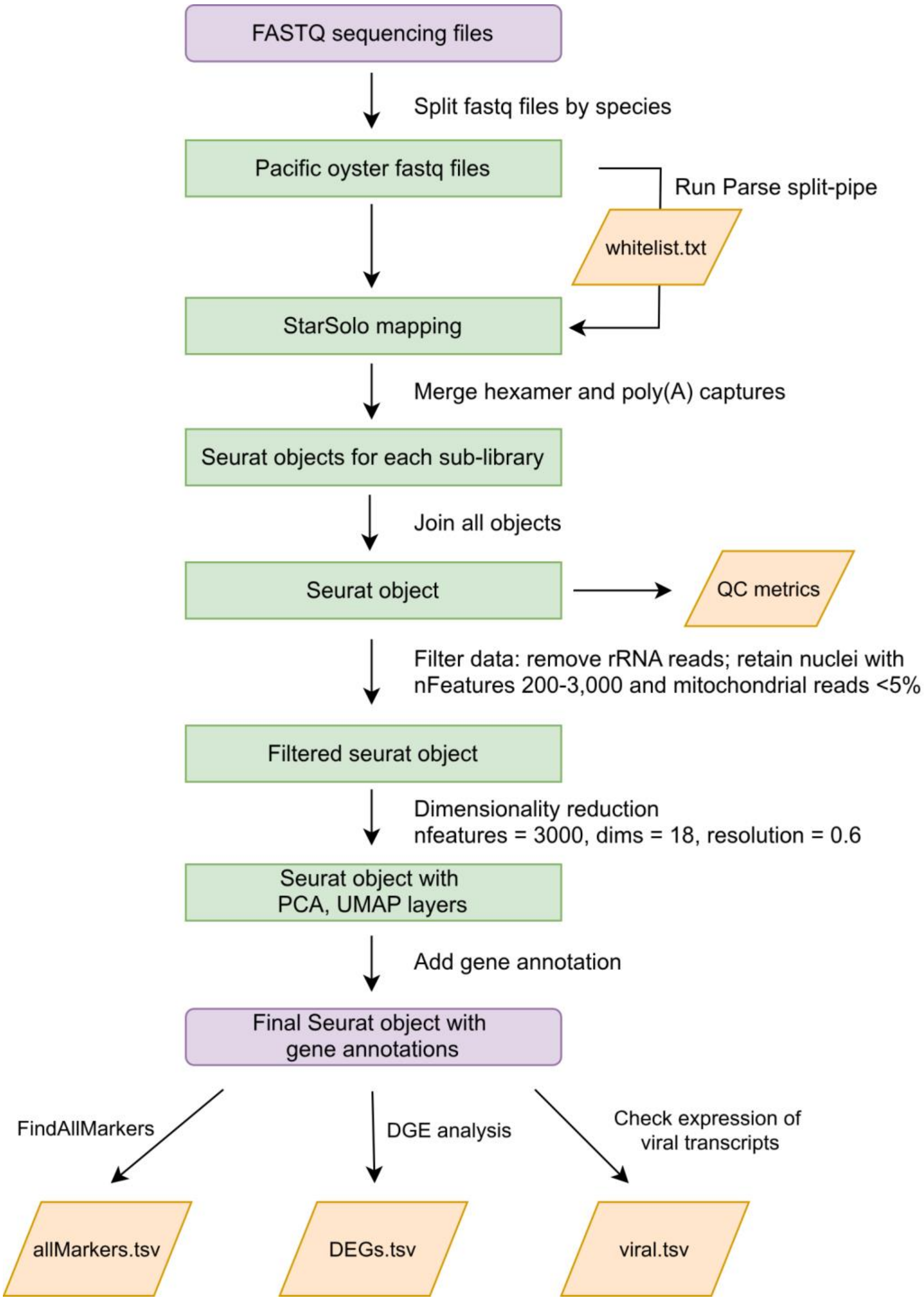


**Figure S1. Bioinformatics workflow for single-nucleus RNA sequencing analysis.**

FASTQ sequencing files were aligned and quantified using STARsolo. Count matrices from eight sub-libraries were imported into Seurat and merged into a single Seurat object. Standard quality-control filtering was applied, and the final dataset was used for marker gene identification, differential gene expression analysis, and evaluation of viral transcript expression.

A

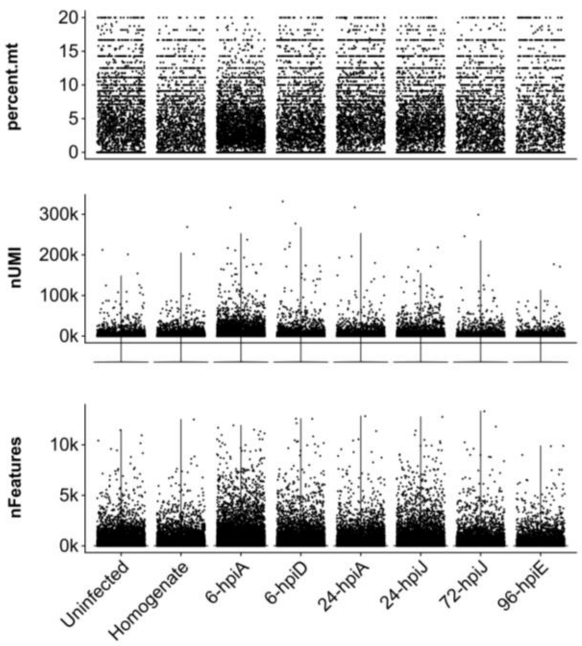


B

| Sample | Number of Nuclei | Total UMIs | Per Nucleus Summary: Mean UMIs | Mean nFeatures | Mean percent.mt |
|---|---|---|---|---|---|
| Uninfected | 147,456 ( 2,603) | 18,348,298 ( 7,449,236) | 124 (2,862) | 35 (813) | 3.1 (1.7) |
| Homogenate-only | 147,456 ( 3,483) | 23,209,046 (13,577,461) | 157 (3,898) | 32 (721) | 2.1 (1.6) |
| 6-hpiA | 147,456 ( 3,898) | 57,711,604 (22,055,463) | 391 (5,658) | 67 (934) | 2.4 (1.6) |
| 6-hpiD | 147,456 ( 3,180) | 26,302,611 (10,054,464) | 178 (3,162) | 43 (848) | 2.6 (1.6) |
| 24-hpiA | 147,456 ( 2,704) | 22,064,804 ( 9,136,217) | 150 (3,379) | 35 (697) | 3.3 (1.9) |
| 24-hpiJ | 147,456 ( 2,975) | 27,336,090 (10,244,751) | 185 (3,444) | 43 (859) | 2.7 (1.4) |
| 72-hpiJ | 147,456 ( 2,830) | 17,438,151 ( 7,750,000) | 118 (2,739) | 31 (695) | 2.3 (1.3) |
| 96-hpiE | 147,456 ( 2,067) | 11,720,784 ( 5,364,829) | 79 (2,595) | 22 (668) | 2.8 (1.9) |
| Total | 1,179,648 (23,740) | 204,131,390 (85,632,420) | 173 (3,607) | 38 (790) | 2.7 (1.6) |

**Figure S2. Quality Control metrics across samples in the merged single-nucleus RNA-sequencing datasets.**

**(A)** Violin plots show mitochondrial read percentage, UMIs per nucleus, and genes detected per nucleus across samples in the merged dataset before filtering. Each dot corresponds to an individual nucleus. To improve visualisation, the mitochondrial read plot was generated from a random subset of 50,000 nuclei. **(B)** Summary of quality-control (QC) metrics before and after filtering. QC metrics are shown for each sample and for the combined dataset; values are presented for the raw merged dataset, with the corresponding filtered values in parentheses. Reported metrics include the number of nuclei, total UMIs, mean UMIs per nucleus, mean genes per nucleus, and mean mitochondrial read percentage. Filtering retained nuclei with 200–3,000 detected genes and <5% mitochondrial UMIs; ribosomal RNA genes were removed from the expression matrix.

**Figure S3 A: gill and mantle group**

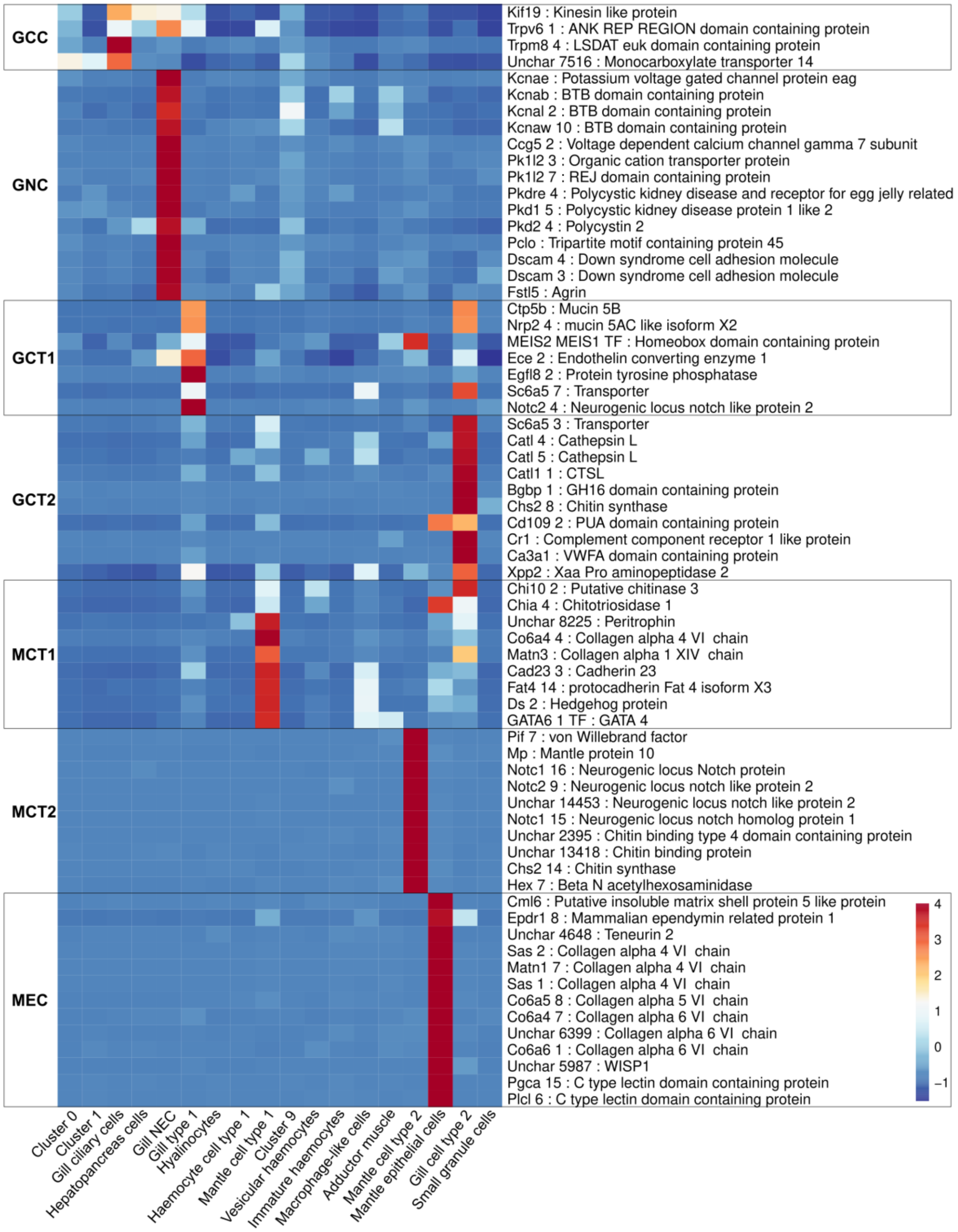

**Figure S3 B: rest of the clusters**

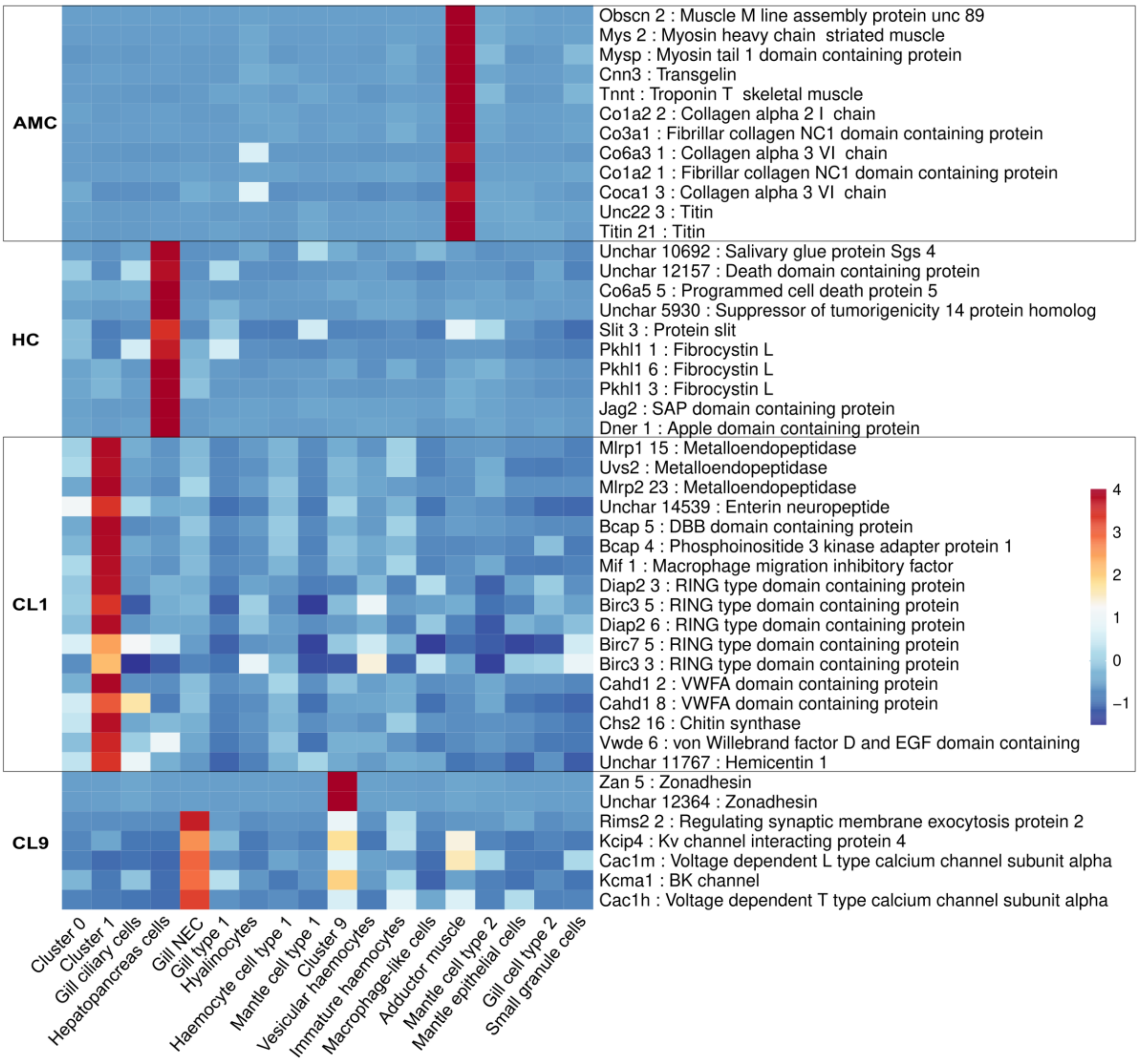

**Figure S3 C: haemocyte group**

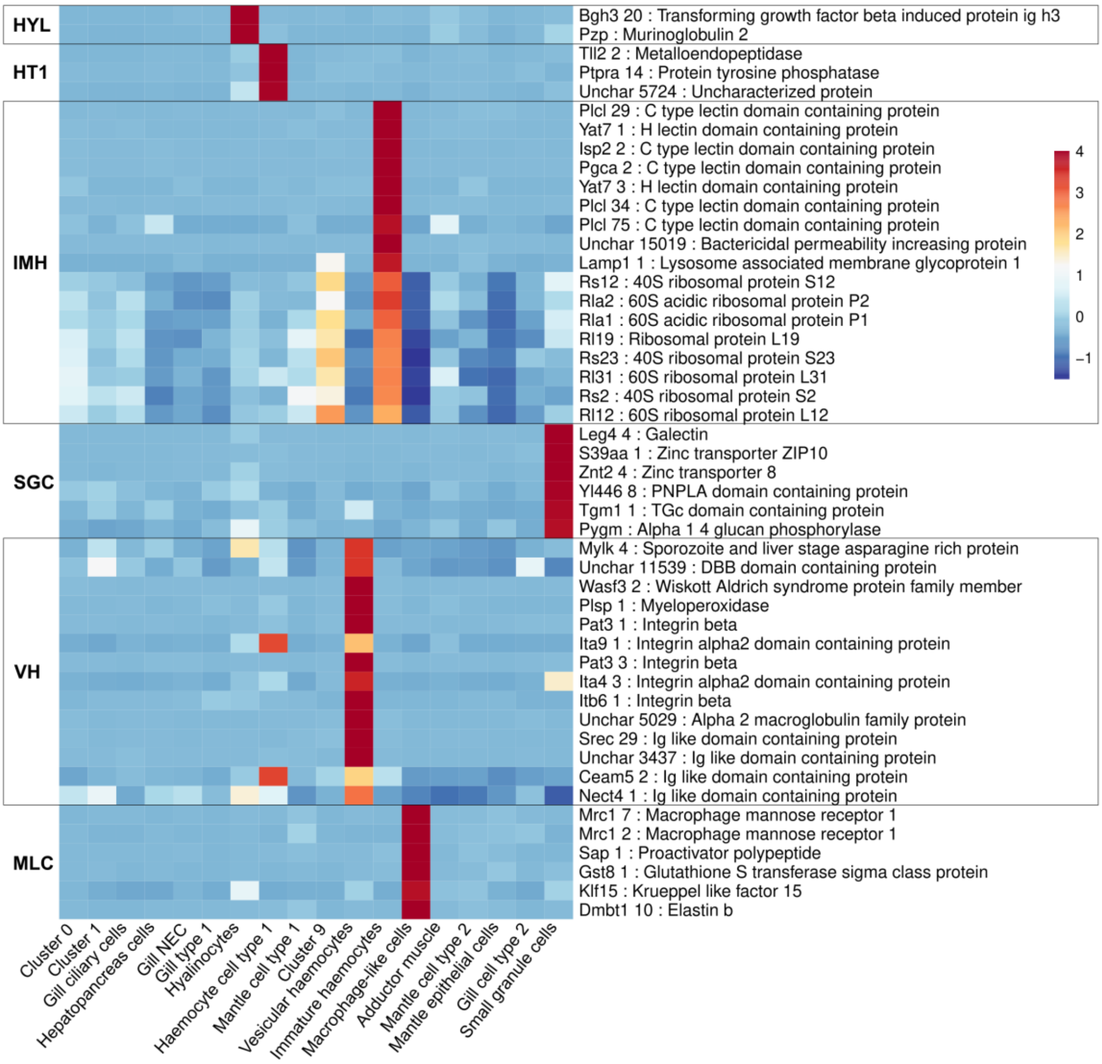


**Figure S3. Expression of key marker genes across transcriptomic clusters described in the Results section.**

Heatmaps depict the average expression of selected marker genes for each transcriptomic cluster. Rows correspond to genes, columns to cell identities, and the colour scale indicates mean gene expression within each cluster. For clarity, clusters are grouped into three categories: **(A)** gill and mantle cell types; **(B)** other cell types, including adductor muscle, hepatopancreas, cluster 1, and cluster 9; and **(C)** haemocytes. Abbreviations shown to the left of each heatmap denote cell identities: GCC, gill ciliary cells; GNC, gill neuroepithelial cells; GCT1, gill cell type 1; GCT2, gill cell type 2; MCT1, mantle cell type 1; MCT2, mantle cell type 2; MEC, mantle epithelial cells; AMC, adductor muscle cells; HC, hepatopancreas cells; CL1, cluster 1; CL9, cluster 9; HYL, hyalinocytes; HT1, haemocyte cell type 1; IMH, immature haemocytes; SGC, small granule cells; VH, vesicular haemocytes; MLC, macrophage-like cells.

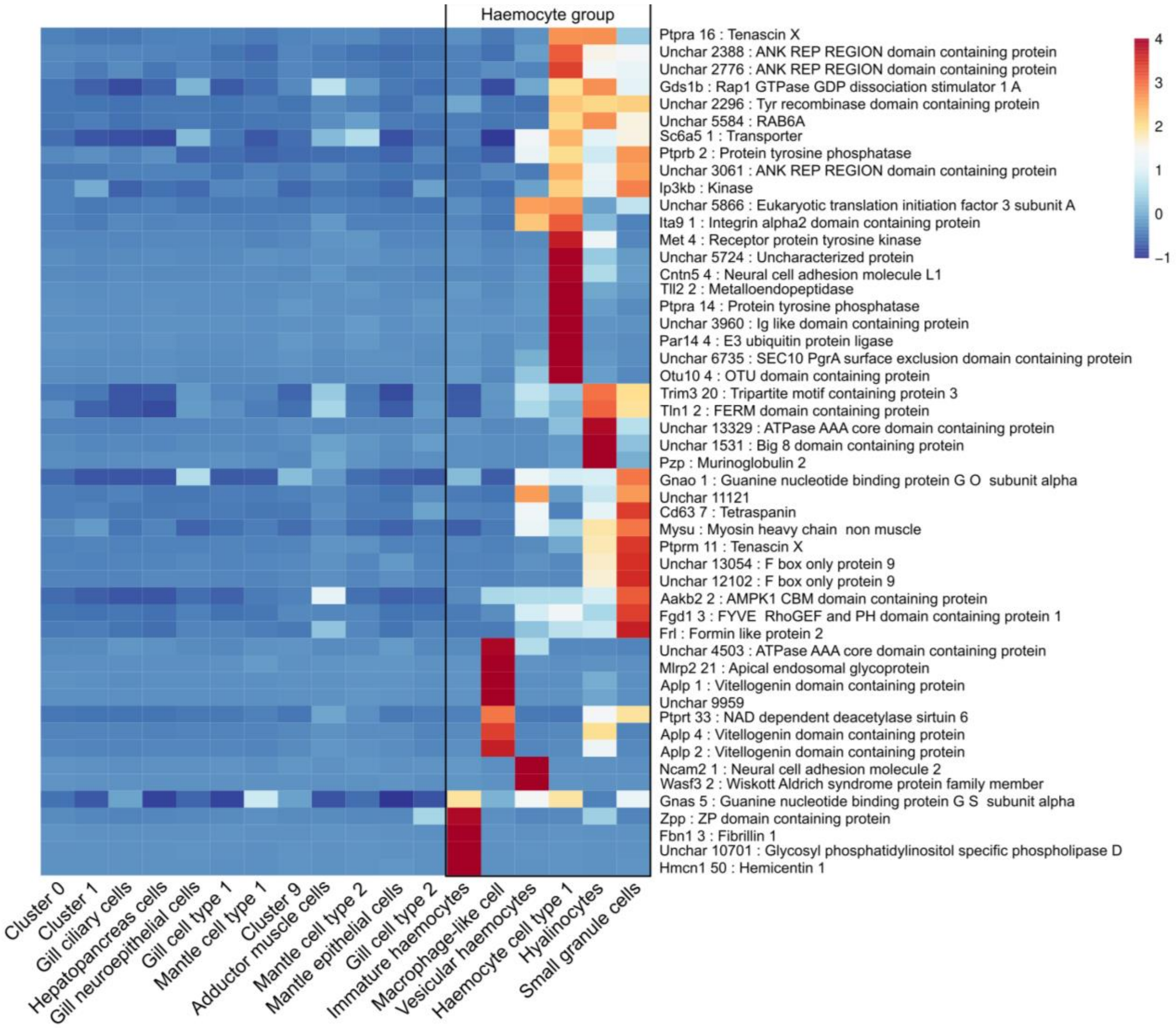


**Figure S4. Conserved marker expression in the haemocyte group.**

Heatmap showing the expression of the top 50 conserved marker genes for the haemocyte group. The haemocyte group comprises the following cell types: immature haemocytes, macrophage-like cells, vesicular haemocytes, haemocyte cell type 1, hyalinocytes, and small granule cells. Conserved markers were identified using the Seurat function *FindConservedMarkers*. The heatmap was generated using *pheatmap* and displays average gene expression values (*AverageExpression*) for the top 50 conserved markers across all transcriptomic clusters. Each row represents a gene, and each column represents a transcriptomic cluster. Clusters belonging to the haemocyte group are highlighted with a rectangular box on the right, and the colour scale representing expression levels is shown alongside the heatmap.

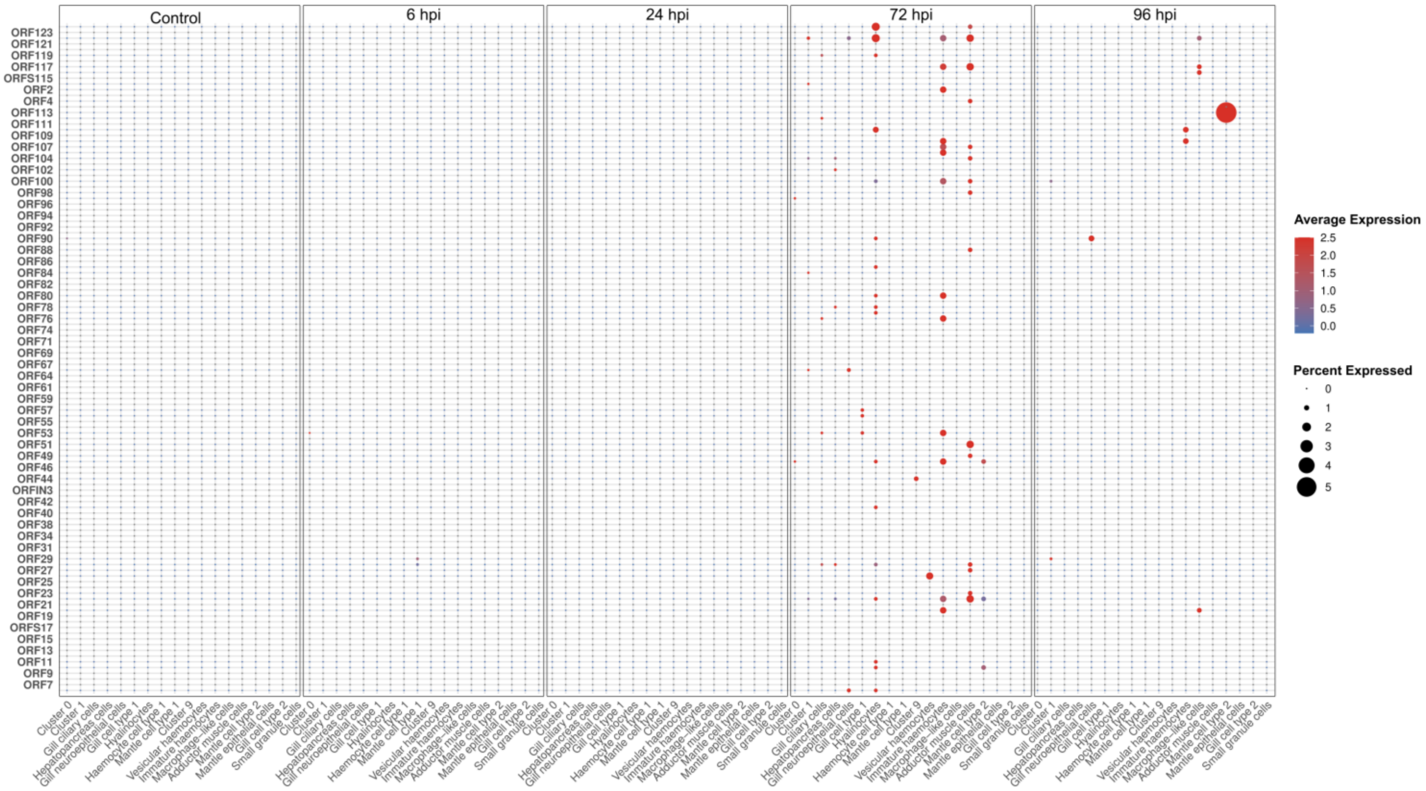


**Figure S5. Expression of OsHV-1 transcripts across samples.**

Dot plot showing the expression of all OsHV-1 transcripts in control and OsHV1-infected Pacific oysters sampled at 6, 24, 72, and 96 hours post infection (hpi). Rows represent OsHV-1 open reading frames (ORFs), and columns correspond to transcriptomic clusters within each sample. Dot colour indicates the average expression level; dot size represents the percentage of nuclei expressing each transcript.

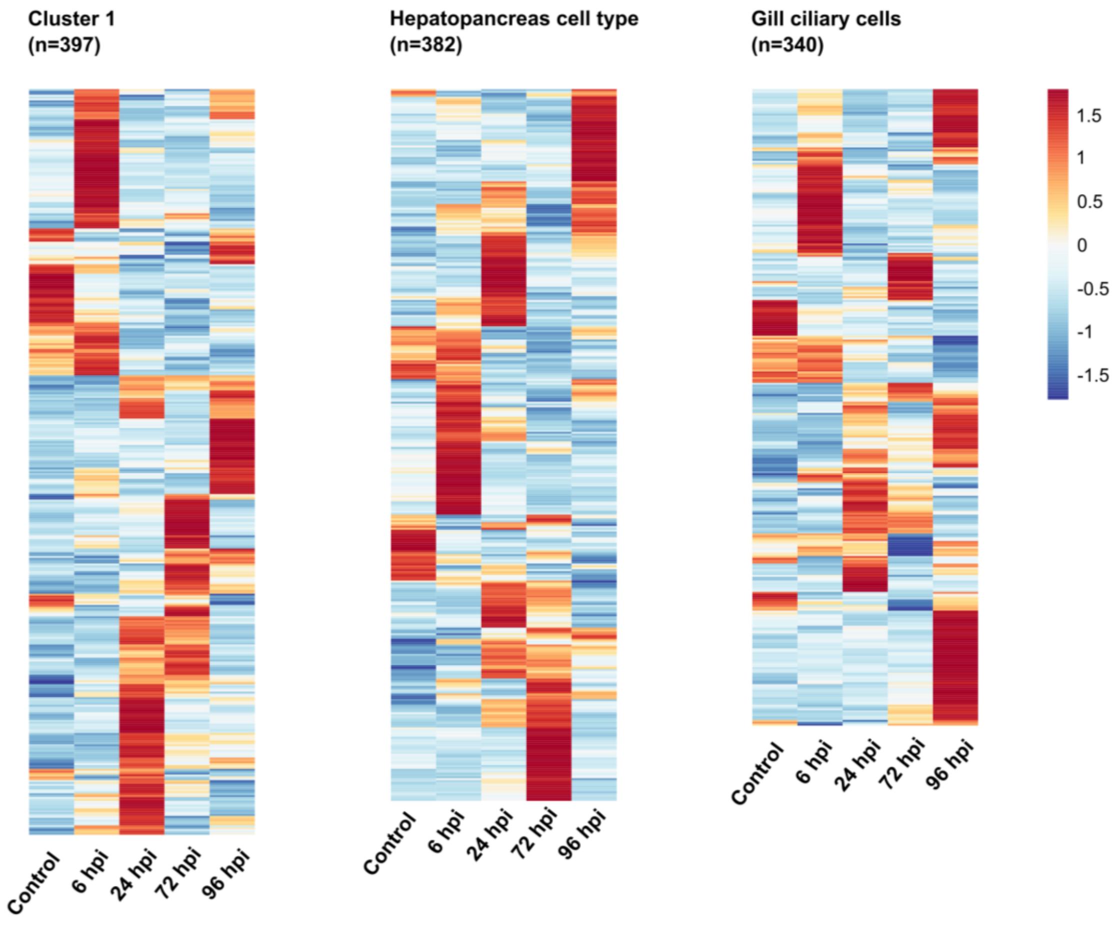


**Figure S6. Heatmaps of differentially expressed genes in clusters with the highest transcriptional response to infection.**

Heatmaps for the three clusters with the highest number of differentially expressed genes (DEGs): cluster 1, hepatopancreas cell type, and gill ciliary cells. For each cluster, DEGs were identified by comparing control versus infected oysters at 6-, 24-, 72-, and 96-hours post-infection (hpi). A unified DEG list for each cluster was generated by combining all DEGs from these pairwise comparisons, and the unique genes (total number shown on the top in parentheses) were used to plot aggregate expression across samples. Each row represents a gene, and each column represents a sample group. The colour scale representing expression levels is displayed on the right.

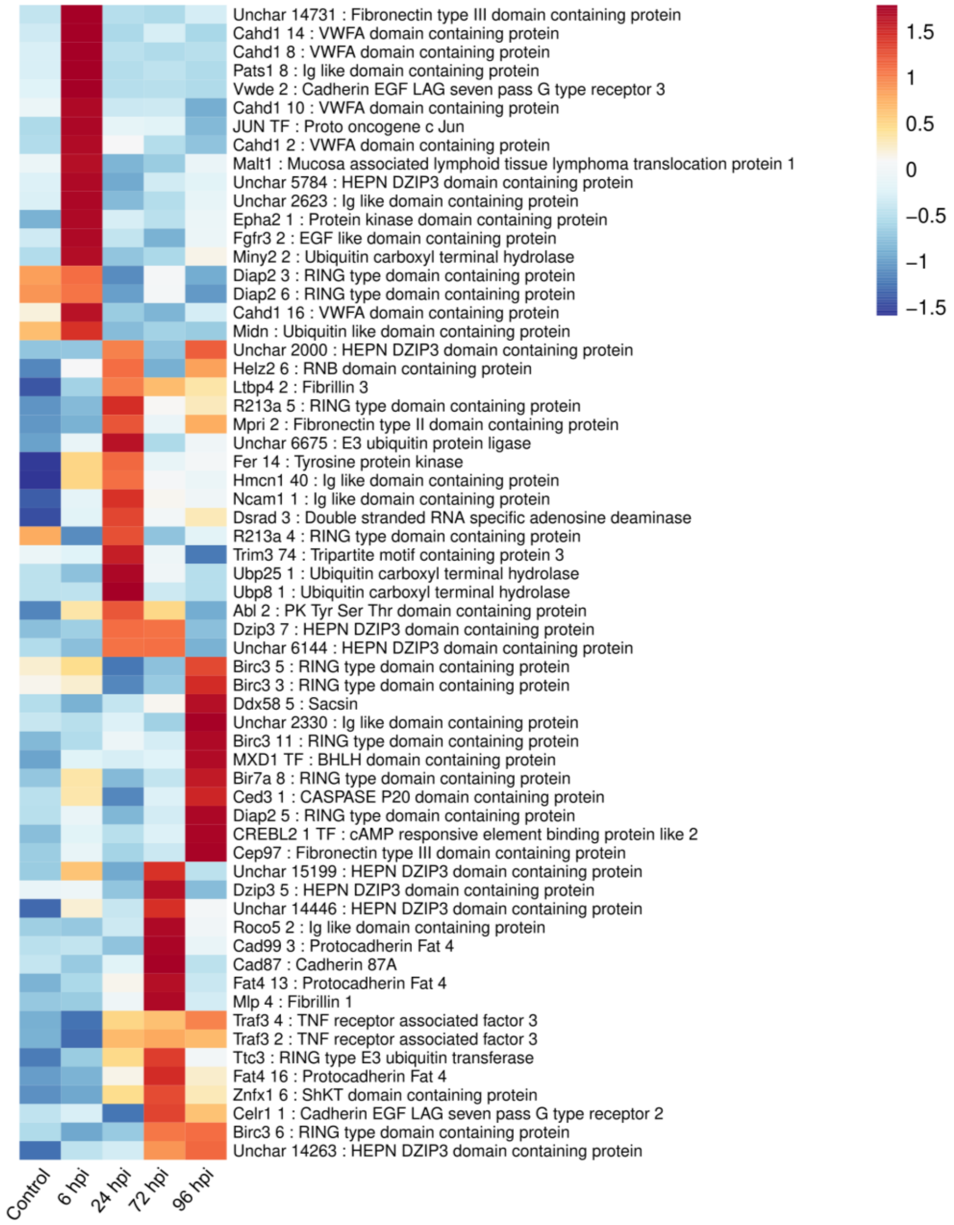


**Figure S7. Immune modulatory genes upregulated in cluster 1 in response to OsHV-1 infection.**

Heatmap showing expression of key immune modulatory genes that were upregulated in response to OsHV-1 infection. These genes were derived from cluster-specific differentially expressed genes using an annotation-guided filtering strategy. Gene descriptions were queried for terms associated with conserved innate immune functions, including RNA sensing (Ddx58, Znfx1, Dsrad), ubiquitin-mediated signalling (TRIM family proteins, E3 ligases), apoptosis (Birc, Ced3), and immune signalling pathways (Traf3, Malt1, Jun). Rows represent gene, columns represent samples. The colour scale representing expression levels is shown on the right.

**Figure S8A**

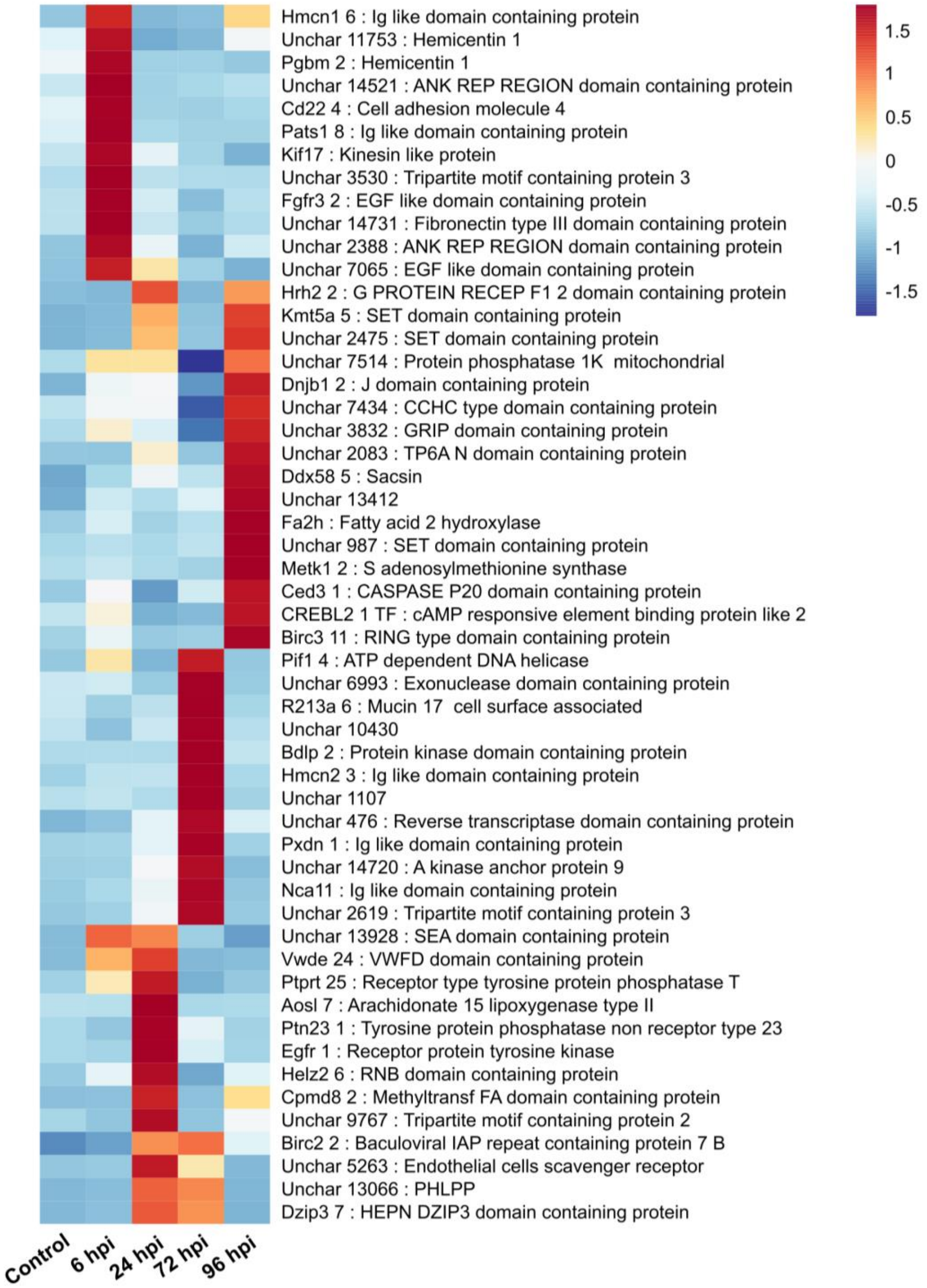

Hmcn1 6 : Ig like domain containing protein
Unchar 11753 : Hemicentin 1
Pgbm 2 : Hemicentin 1
Unchar 14521 : ANK REP REGION domain containing protein
Cd22 4 : Cell adhesion molecule 4
Pats1 8 : Ig like domain containing protein
Kif17 : Kinesin like protein
Unchar 3530 : Tripartite motif containing protein 3
Fgfr3 2 : EGF like domain containing protein
Unchar 14731 : Fibronectin type III domain containing protein
Unchar 2388 : ANK REP REGION domain containing protein
Unchar 7065 : EGF like domain containing protein
Hrh2 2 : G PROTEIN RECEP F1 2 domain containing protein
Kmt5a 5 : SET domain containing protein
Unchar 2475 : SET domain containing protein
Unchar 7514 : Protein phosphatase 1K mitochondrial
Dnjb1 2 : J domain containing protein
Unchar 7434 : CCHC type domain containing protein
Unchar 3832 : GRIP domain containing protein
Unchar 2083 : TP6A N domain containing protein
Ddx58 5 : Sacsin
Unchar 13412
Fa2h : Fatty acid 2 hydroxylase
Unchar 987 : SET domain containing protein
Metk1 2 : S adenosylmethionine synthase
Ced3 1 : CASPASE P20 domain containing protein
CREBL2 1 TF : cAMP responsive element binding protein like 2
Birc3 11 : RING type domain containing protein
Pif1 4 : ATP dependent DNA helicase
Unchar 6993 : Exonuclease domain containing protein
R213a 6 : Mucin 17 cell surface associated
Unchar 10430
Bdlp 2 : Protein kinase domain containing protein
Hmcn2 3 : Ig like domain containing protein
Unchar 1107
Unchar 476 : Reverse transcriptase domain containing protein
Pxdn 1 : Ig like domain containing protein
Unchar 14720 : A kinase anchor protein 9
Nca11 : Ig like domain containing protein
Unchar 2619 : Tripartite motif containing protein 3
Unchar 13928 : SEA domain containing protein
Vwde 24 : VWFD domain containing protein
Ptprt 25 : Receptor type tyrosine protein phosphatase T
Aosl 7 : Arachidonate 15 lipoxygenase type II
Ptn23 1 : Tyrosine protein phosphatase non receptor type 23
Egfr 1 : Receptor protein tyrosine kinase
Helz2 6 : RNB domain containing protein
Cpmd8 2 : Methyltransf FA domain containing protein
Unchar 9767 : Tripartite motif containing protein 2
Birc2 2 : Baculoviral IAP repeat containing protein 7 B
Unchar 5263 : Endothelial cells scavenger receptor
Unchar 13066 : PHLPP
Dzip3 7 : HEPN DZIP3 domain containing protein
1.5
1
0.5
0
-0.5
-1
-1.5
Control
6 hpi
24 hpi
72 hpi
96 hpi

**Figure S8B**

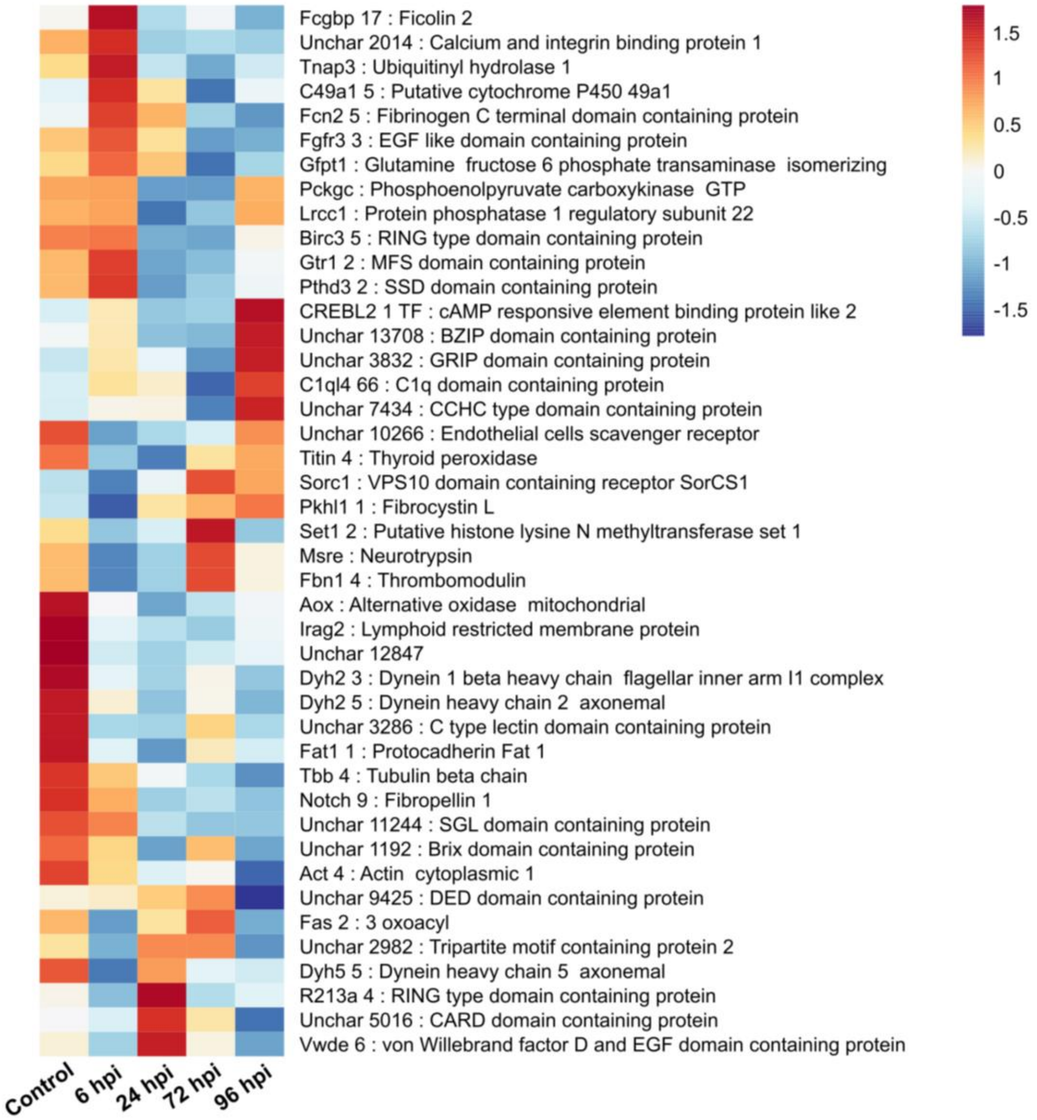


**Figure S8: Top genes differentially expressed in the cluster hepatopancreas cell type.**

Heatmap representing the expression of the 15 most significant DEGs per comparison identified in hepatopancreas cells for each of the four pairwise control vs infected stages (6, 24, 72, 96 hpi). All five conditions are included to visualize changes over time. Genes are arranged by row, samples by column. Significantly upregulated **(A)** and downregulated **(B)** genes are displayed in separate heatmaps, colour scale representing expression levels is shown on the right.

**Figure S9A**

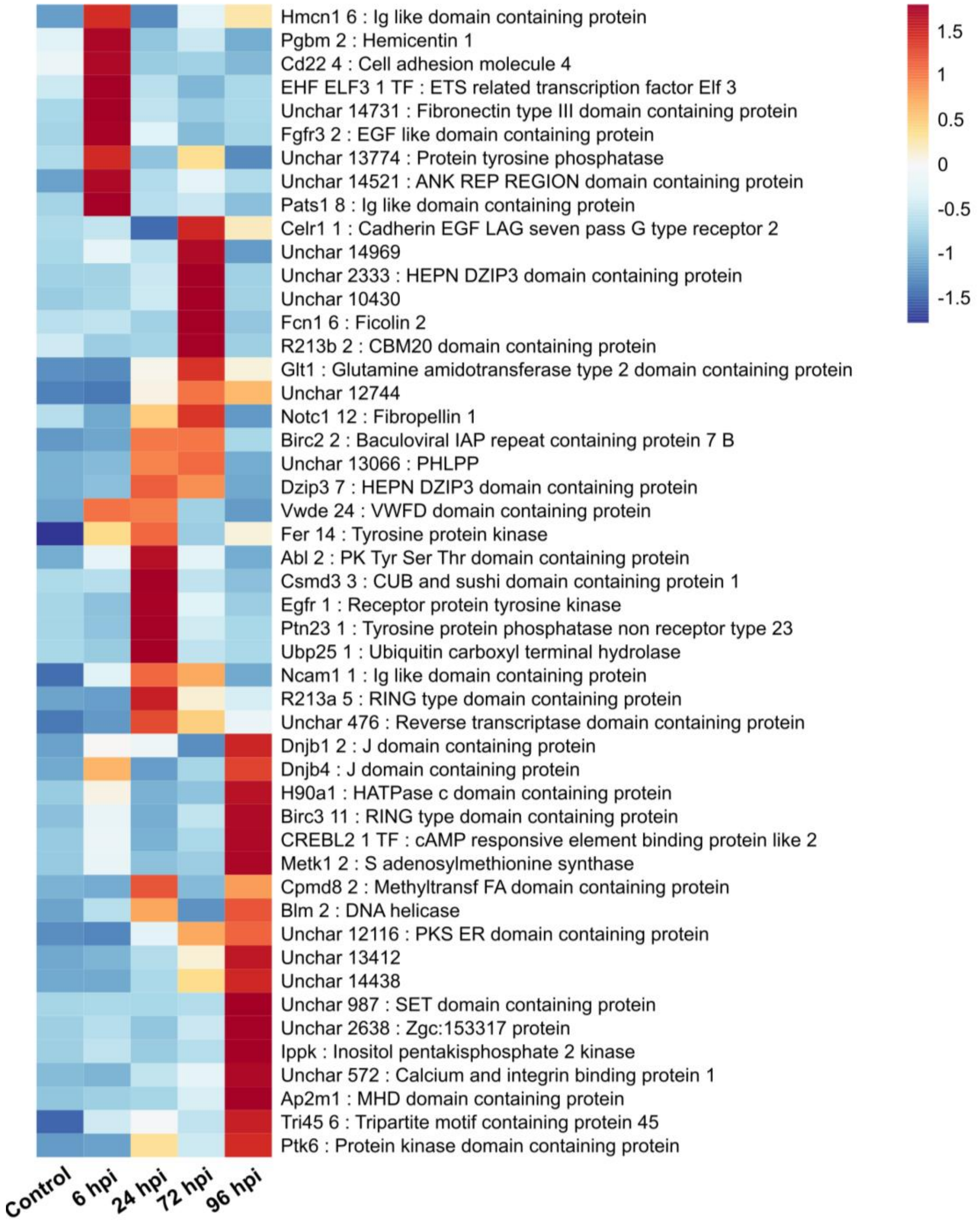
Hmcn1 6 : Ig like domain containing protein
Pgbm 2 : Hemicentin 1
Cd22 4 : Cell adhesion molecule 4
EHF ELF3 1 TF : ETS related transcription factor Elf 3
Unchar 14731 : Fibronectin type III domain containing protein
Fgfr3 2 : EGF like domain containing protein
Unchar 13774 : Protein tyrosine phosphatase
Unchar 14521 : ANK REP REGION domain containing protein
Pats1 8 : Ig like domain containing protein
Celr1 1 : Cadherin EGF LAG seven pass G type receptor 2
Unchar 14969
Unchar 2333 : HEPN DZIP3 domain containing protein
Unchar 10430
Fcn1 6 : Ficolin 2
R213b 2 : CBM20 domain containing protein
Glt1 : Glutamine amidotransferase type 2 domain containing protein
Unchar 12744
Notc1 12 : Fibropellin 1
Birc2 2 : Baculoviral IAP repeat containing protein 7 B
Unchar 13066 : PHLPP
Dzip3 7 : HEPN DZIP3 domain containing protein
Vwde 24 : VWFD domain containing protein
Fer 14 : Tyrosine protein kinase
Abl 2 : PK Tyr Ser Thr domain containing protein
Csmd3 3 : CUB and sushi domain containing protein 1
Egfr 1 : Receptor protein tyrosine kinase
Ptn23 1 : Tyrosine protein phosphatase non receptor type 23
Ubp25 1 : Ubiquitin carboxyl terminal hydrolase
Ncam1 1 : Ig like domain containing protein
R213a 5 : RING type domain containing protein
Unchar 476 : Reverse transcriptase domain containing protein
Dnjb1 2 : J domain containing protein
Dnjb4 : J domain containing protein
H90a1 : HATPase c domain containing protein
Birc3 11 : RING type domain containing protein
CREBL2 1 TF : cAMP responsive element binding protein like 2
Metk1 2 : S adenosylmethionine synthase
Cpmd8 2 : Methyltransf FA domain containing protein
Blm 2 : DNA helicase
Unchar 12116 : PKS ER domain containing protein
Unchar 13412
Unchar 14438
Unchar 987 : SET domain containing protein
Unchar 2638 : Zgc:153317 protein
Ippk : Inositol pentakisphosphate 2 kinase
Unchar 572 : Calcium and integrin binding protein 1
Ap2m1 : MHD domain containing protein
Tri45 6 : Tripartite motif containing protein 45
Ptk6 : Protein kinase domain containing protein
1.5
1
0.5
0
-0.5
-1
-1.5
Control
6 hpi
24 hpi
72 hpi
96 hpi

Figure S9B

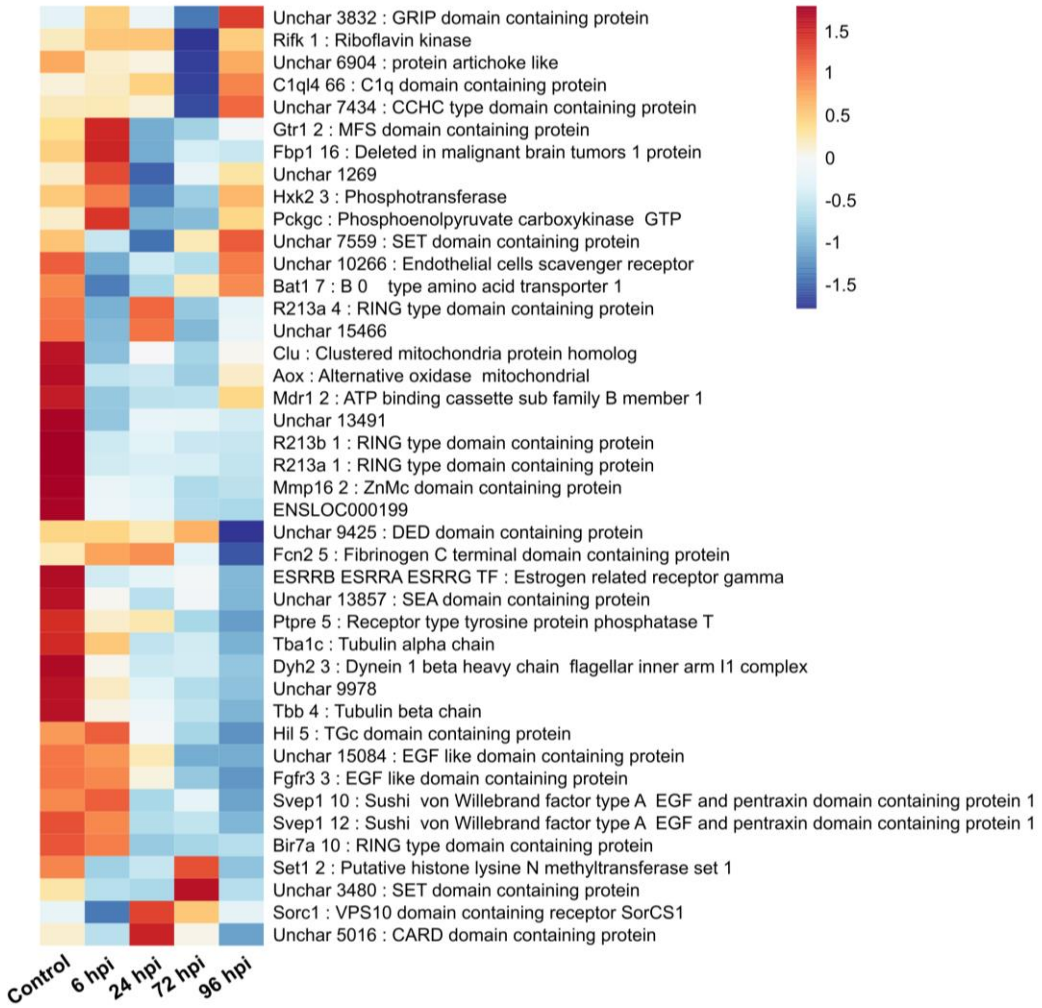


**Figure S9: Top differentially expressed genes identified in the gill ciliary cell cluster.**

Heatmap illustrating the expression of the top 15 DEGs in gill ciliary cells across the four pairwise comparisons between control and infected oysters at 6, 24, 72, and 96 hpi. Expression patterns for all five conditions are plotted to highlight temporal dynamics. Each row shows a Pacific oyster gene, and each column corresponds to a sample. Separate panels depict genes upregulated **(A)** or downregulated **(B)** upon infection. The colour scale for expression is shown on the right.

| Sample subtype | Total nuclei | Viral-positive nuclei (n, %) |
|---|---|---|
| Adductor muscle cells_72hpi | 123 | 11 (8.9%) |
| Immature haemocytes_72hpi | 74 | 6 (8.1%) |
| Hyalinocytes_72hpi | 166 | 11 (6.6%) |
| Mantle cell type 2_96hpi | 19 | 1 (5.3%) |
| Mantle cell type 2_72hpi | 115 | 3 (2.6%) |
| Immature haemocytes_96hpi | 88 | 2 (2.3%) |
| Gill neuroepithelial cells_72hpi | 151 | 3 (2%) |
| Hepatopancreas cells_72hpi | 278 | 5 (1.8%) |
| Gill ciliary cells_72hpi | 295 | 5 (1.7%) |
| Vesicular haemocytes_72hpi | 65 | 1 (1.5%) |
| Cluster 1_72hpi | 383 | 5 (1.3%) |
| Gill neuroepithelial cells_96hpi | 85 | 1 (1.2%) |

**Table S1. Distribution of nuclei expressing viral transcripts across cell subtypes and infection conditions**

Table summarising the number and proportion of single nuclei expressing viral transcripts for each annotated cell identity and infection condition. Cell identities were defined by combining cluster annotation with infection timepoint (control, 6 hpi, 24 hpi, 72 hpi, 96 hpi) to generate sample subtype. Viral-positive nuclei were defined as those containing at least one UMI mapping to viral genes (ORFs). Only sample subtypes with >1% viral-positive nuclei are shown.
